# Widespread promiscuous alkaline phosphatases underscore early microbial phosphite utilization

**DOI:** 10.1101/2025.04.29.651336

**Authors:** Morito Sakuma, Naoki Konno, Sevan Gholipour, John Z Chen, Nobuhiko Tokuriki

## Abstract

Phosphate is often a limiting resource, directly affecting the availability of key biomolecules such as nucleotides. To cope with phosphate scarcity, bacteria have evolved enzymes that utilize alternative phosphorus compounds, including phosphite (Pt). Although a few enzymes oxidize Pt to produce phosphate, the enzymes responsible for Pt oxidation in many environmental bacteria remain unidentified, and the role of microbial Pt oxidation in the global phosphorus cycle is not yet fully understood. In this study, we performed bioinformatic analyses of three Pt-oxidizing enzymes: the native Pt oxidase phosphite dehydrogenase (PtxD), and two promiscuous Pt oxidases, alkaline phosphatase (AP) and carbon-phosphorus (CP) lyase. Among these, AP was found to be widely distributed across bacteria since the early stages of their evolution. In contrast, PtxD emerged later in a limited number of bacterial lineages that had lost AP. Our biochemical characterizations revealed that most extant and reconstructed ancestral APs tested exhibited Pt oxidation activity. Moreover, disruption of active-site residues diminished Pt oxidase activity in AP, while only partially affecting its native function. This promiscuous function of AP reveals an overlooked mechanism in bacterial phosphate metabolism and underscores the role of Pt in the cycling of bioavailable phosphorus in ecosystems.

## Introduction

Phosphate is essential for life, forming key components such as ATP, nucleic acids, and cell membranes. Nevertheless, phosphate is often limited in many ecosystems, directly constraining the growth of organisms [1-3]. To overcome this limitation, microbes utilize alternative sources of dissolved organic phosphorus (DOP) via enzymes such as phosphatases [3-7]. Phosphonates are also abundant and serve as a source of phosphate through the cleavage of carbon-phosphorus (C-P) bonds by CP lyases and phosphonatases [1-3, 8, 9]. In addition to these organic sources, phosphite (Pt), a reduced and highly soluble inorganic phosphorus compound, is of particular interest due to its ecological role in the global phosphorus cycle and microbial metabolism [1-3, 10]. Pt is thought to have been prevalent in early Earth ecosystems, especially under reducing environments before the Great Oxidation Event (GOE) (> 2.5 Ga) [1,3,11-13]. Its abundance subsequently declined after the GOE (< 2.3 Ga), due to a reduction in Pt-generating processes and shifts in Earth’s redox conditions. However, Pt is still frequently detected up to micromolar concentrations across diverse modern environments, including soil, aquatic systems, and geothermal settings [1-3, 14], highlighting its ongoing ecological relevance.

Pt is a kinetically stable compound, requiring an activation energy of 370 kJ to break its P–H bond [11]. Therefore, microbes rely on specialized enzymes—such as Pt oxidases—to convert Pt into phosphate. To date, however, only one enzyme family, phosphite dehydrogenase (PtxD), has been classified as catalyzing Pt oxidation as its primary function [15, 16]. PtxD efficiently catalyzes Pt oxidation (*k*_cat_/*K*_M_ ∼ 10^5^ M^-1^s^-1^) using NAD^+^ as a cofactor and has been identified as a part of the *ptx* operon (**Fig. 1A**) [3, 17]. In addition to generating phosphate from Pt to support bacterial growth, PtxD is also capable of utilizing phosphite as an exclusive electron donor, facilitating the coupling of Pt oxidation to the reduction of carbon dioxide, sulfate, or nitrate in the process known as dissimilatory phosphite oxidation (DPO) [3, 18]. Despite its efficiency, PtxD is surprisingly rare in bacterial genomes, primarily identified in species such as *Pseudomonas stutzeri* (*P. stutzeri*) and certain anaerobes [3, 19, 20]. A recent phylogenetic study by Boden *et al*. also suggested that *ptxD* (encoding PtxD) genes underwent significant expansion only after the GOE [21]. These observations raise the question of whether PtxD is the sole enzyme responsible for Pt oxidation in nature, and suggest that global microbially driven Pt oxidation may depend on only a limited number of bacterial species harboring PtxD. However, studies have shown that a substantial proportion (10–67%) of bacterial isolates can grow in media containing Pt as the sole phosphorus source, suggesting that other, yet unidentified, enzymes may also be involved in Pt oxidation [22].

**Fig. 1.**
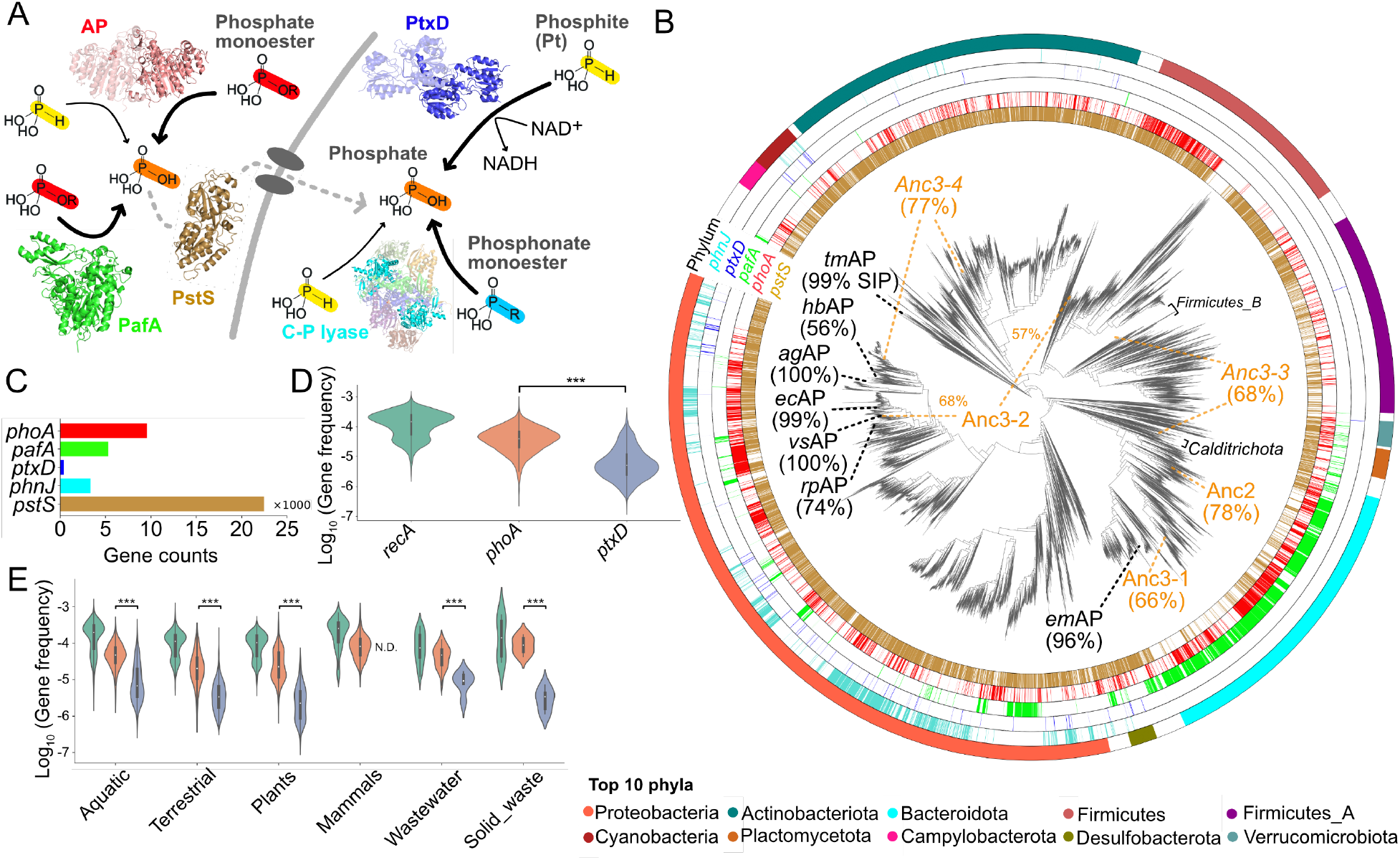
Distribution of phosphite (Pt) oxidase genes on the phylogenetic tree of bacterial genomes. (A) Phosphate production and transportation in bacteria. Alkaline phosphatase (AP, PDB: 1ALK), located in the periplasm, catalyzes the hydrolysis of phosphate monoester substrates and the Pt oxidation reaction to produce phosphate. PafA (PDB: 5TJ3), another member of the AP superfamily, also catalyzes the hydrolysis of phosphate monoesters. The phosphate-binding protein PstS (PDB: 1A40) and *pho* regulon proteins transport phosphate into the cytoplasm. PtxD (PDB: 4E5K) catalyzes the Pt oxidation reaction in the cytoplasm using NAD^+^ as a cofactor. Carbon-phosphorus (CP) lyase (PDB: 4XB6) degrades phosphonate monoester substrates to produce phosphate and catalyzes Pt oxidation. Native and promiscuous functions are indicated by thick and thin arrows, respectively. (B) Distribution of the *phoA* (red), *pafA* (green), *ptxD* (blue), and *phnJ* (cyan) genes mapped onto the bacterial phylogenetic tree. The presence of each gene is indicated by lines on the circle. Bacterial phyla are also mapped onto the tree. The positions of the twelve *phoA* genes exhibiting Pt oxidation activity are shown, along with the *phoA* genes in bacterial genomes that have the highest sequence identity percentage (SIP) to them. SIP was calculated using BLASTp. (C) The abundance of enzyme genes across all bacterial phyla. (D), (E) Metagenomic analysis of the *phoA* and *ptxD* genes. (D) Violin plots showing the relative frequencies of the *recA, phoA*, and *ptxD* genes across all environments. The *recA* gene was used as a control [30]. *** indicates a p-value < 0.001 in the Kolmogorov-Smirnov (KS) test. (E) Relative frequencies of the *recA, phoA*, and *ptxD* genes in each environment. *N*.*D*. indicates that no *ptxD* genes were detected in animal-associated environments.

Two other enzymes, CP lyase (encoded by the *phn* genes) and alkaline phosphatase (encoded by the *phoA* gene), have been reported to exhibit Pt oxidation activity as a non-primary or promiscuous function. While CP lyase and its *phn* operon in *Escherichia coli* (*E. coli*) and *P. stutzeri* facilitate phosphate assimilation from phosphonate substrates [1, 8], they can also contribute to Pt oxidation (**Fig. 1A**) [1, 23]. However, the *phn* genes also show limited distribution among environmental microbes and are believed to have expanded after the GOE [21]. AP is a widely distributed periplasmic phosphomonoesterase (PMEase) in bacteria that produces phosphate from phosphoester compounds [3, 4] (**Fig. 1A**). APs often catalyze multiple promiscuous reactions, including Pt oxidation [24–27].

Interestingly, although both AP and CP lyase exhibit significantly lower Pt oxidation activity than PtxD—with AP showing a turnover rate nearly 10,000-fold lower—it has been demonstrated that heterologous expression of *E. coli* AP and *P. stutzeri* CP lyase enables bacterial growth in Pt-containing media [24, 28]. These observations raise a pivotal question: Do these promiscuous enzymes play a physiological role in microbial Pt oxidation, or are they merely serendipitous, non-physiological promiscuous activities? A comprehensive understanding of the distributions of PtxD and promiscuous Pt oxidases among microbes is essential for elucidating the role of microbial Pt oxidation in global phosphorus cycling.

In this study, we address these questions through comprehensive bioinformatic analyses and biochemical characterizations. We found that the *ptxD* gene is exceedingly rare, present in only a few specific bacterial taxa, whereas the *phoA* gene was the most abundant and widely distributed among diverse bacterial phyla. Through experimental characterization of twelve APs, including reconstructed ancestral APs, we demonstrated that Pt oxidase activity is far more widespread among bacterial APs than previously recognized. We propose that PtxD, a specialized Pt oxidase, likely evolved later to confer a growth advantage in Pt-rich environments. In contrast, multifunctional APs may represent the most ancient Pt oxidase, playing a significant role in Pt oxidation from the pre-GOE to the modern era. Finally, we discuss the potential contribution of phosphatase-mediated Pt oxidation to global phosphorus cycling.

## Results

### The distribution of the *ptxD, phnJ, pafA*, and *phoA* genes among bacteria

We first curated and classified homologous sequences for three enzyme families known to catalyze Pt oxidation (**Fig. S1** and see **Methods**)—PtxD (encoded by the *ptxD* gene), CP lyase (*phnJ* gene, which encodes the catalytic subunit (PhnJ) of the CP lyase complex), and AP (*phoA* gene)— as well as PafA (*pafA* gene), an efficient PMEase homologous to AP. We analyzed their distribution across a previously constructed bacterial phylogenetic tree comprising 32,225 representative bacterial species (one genome per species) across 105 phyla [29] (**Fig. 1B**). While the *ptxD* gene is the only native Pt oxidase gene, its orthologs were exceedingly rare, found in only 1.2% of bacterial genomes (373 genomes), mostly within *Proteobacteria* and to a lesser extent in *Actinobacteria* and *Cyanobacteria* (**Fig. 1B-C**). While the *phnJ* gene was more abundant (10%, 3,346 genes) than the *ptxD* gene, it was primarily distributed in *Proteobacteria* (**Fig. 1B-C**). In contrast, the *phoA* gene was widely distributed, identified in approximately 30% of the bacterial genome (9,588 genomes) across various phyla (**Fig. 1B-C**). The broad distribution of the *phoA* gene is especially notable compared to the *pafA* gene, which was found in only 16% of bacterial genomes (5,277 genes) and was predominantly distributed in *Bacteroidota* (69% of the *pafA* genes, 3,626 genes) (**Fig. 1B-C**). Overall, the *phoA*/*pafA* gene pair was present in about 40% of bacterial genomes (**Fig. 1B-C**). To investigate the distribution of the *ptxD* and *phoA* genes among environmental microbes, we searched for their homologs in a large set of metagenomic data [30]. We compared the abundance of the *ptxD* and *phoA* genes with that of the *recA* gene, a common single-copy bacterial marker gene. Both genes exhibited similar frequency to the genomic data; the *phoA* gene was found in about 30% of the *recA* gene, and the *ptxD* gene appeared only in less than 5% (**Fig. 1D**). Moreover, both genes are found in diverse environments; however, the *ptxD* gene was not observed in animal-associated environments (**Fig. 1E**). We also examined the distribution of the phosphate-binding protein PstS (*pstS* gene) (**Fig. 1A**), which shares the same regulon as the *phoA* gene [31] and is considered to have existed since the Archean Eon [21]. Indeed, the *pstS* gene was prevalent among bacterial genomes (22,463 genes, 70% of total genomes) (**Fig. 1B-C**). Therefore, the *phoA* gene, which has also been inferred to be part of the gene repertoire of the last universal common ancestor (LUCA) [32], may have been present in bacteria since ancient times, possibly as early as the Archean Eon, as a member of the same *pho* regulon. We further analyzed the co-occurrence and mutual exclusivity of the *ptxD, phoA*, and *phnJ* genes in the phylogenetic tree. Interestingly, while the *phoA* and *phnJ* genes appeared to be partially co-distributed, the *phoA* and *ptxD* genes exhibited a complementary distribution pattern (**Fig. 1B**). Co-occurrence analysis using evolCCM [33] revealed no significant co-occurrence between the *phoA* and *phnJ* genes (p < 0.005), whereas the *phoA* and *ptxD* genes showed a significantly negative co-occurrence (p < 0.001) (**Fig. S2**), suggesting possible functional redundancy or conflict between the genes. Notably, the *pafA* gene also exhibited significant negative co-occurrence with the *ptxD* gene (p < 0.001), suggesting that the *ptxD* gene may have a function inversely related to phosphatase-associated activity (**Fig. S2**). To further explore the patterns of gene gain, loss, and coexistence during the evolutionary process, we used PastML, which predicts the presence or absence of genes at each ancestral node in the phylogenetic tree [29, 34] (**Fig. S3**). While the *phoA* and *phnJ* genes exhibited co-occurrence in their ancestral states, the *phoA* and *ptxD* genes showed minimal overlap in their ancestral distribution (**Fig. S3**). Interestingly, the gain of the *ptxD* gene was largely restricted to lineages in which the *phoA* gene had been lost (**Fig. S3A**), suggesting that the *ptxD* gene may have provided a similar function to the *phoA* gene.

### Sequence diversity of APs

As the *phoA* genes were the most widely distributed among phosphite (Pt) oxidases, we next explored the diversity and phylogenetic distribution within the AP superfamily (**Fig. 2A**). We collected sequences belonging to the AP superfamily (total of 32,843 sequences), which includes various enzyme families such as phosphodiesterases, sulfatases, PafA, and mutases. Using Sequence similarity networks (SSNs), we isolated a sequence cluster that contained APs (15,722 sequences) (**Fig. S1A**), identified by conserved active site structures including the bimetallic binding site, catalytic residues, and nucleophilic serine, as well as annotated AP sequences (**Fig. S4–6**). Within the AP family, we identified six major subclusters (AP1–6), among which AP3 and AP4 were further subdivided into four (AP3-1 to AP3-4) and two (AP4-1 and AP4-2) subgroups, respectively (**Fig. 2A**). We also generated a phylogenetic tree of the AP family using representative sequences from each subcluster (total of 3,091 sequences) (**Fig. 2B**). The phyla of the host organisms were mapped onto both the SSNs and the phylogenetic tree, color-coded by phylum (**Fig. 2A-B**). Many characterized bacterial and fungal APs, including *Bacillus subtilis* (*bs*AP), *Escherichia coli* (*ec*AP), *Thermotoga maritima* (*tm*AP), and *Saccharomyces cerevisiae* (*sc*AP), were found in AP3 (**Fig. 2A** and **Fig. S6**). All characterized mammalian APs belonged to AP4, while the cold-active *Vibrio sp*. AP (*vs*AP) was located in AP6 [35].

**Fig. 2.**
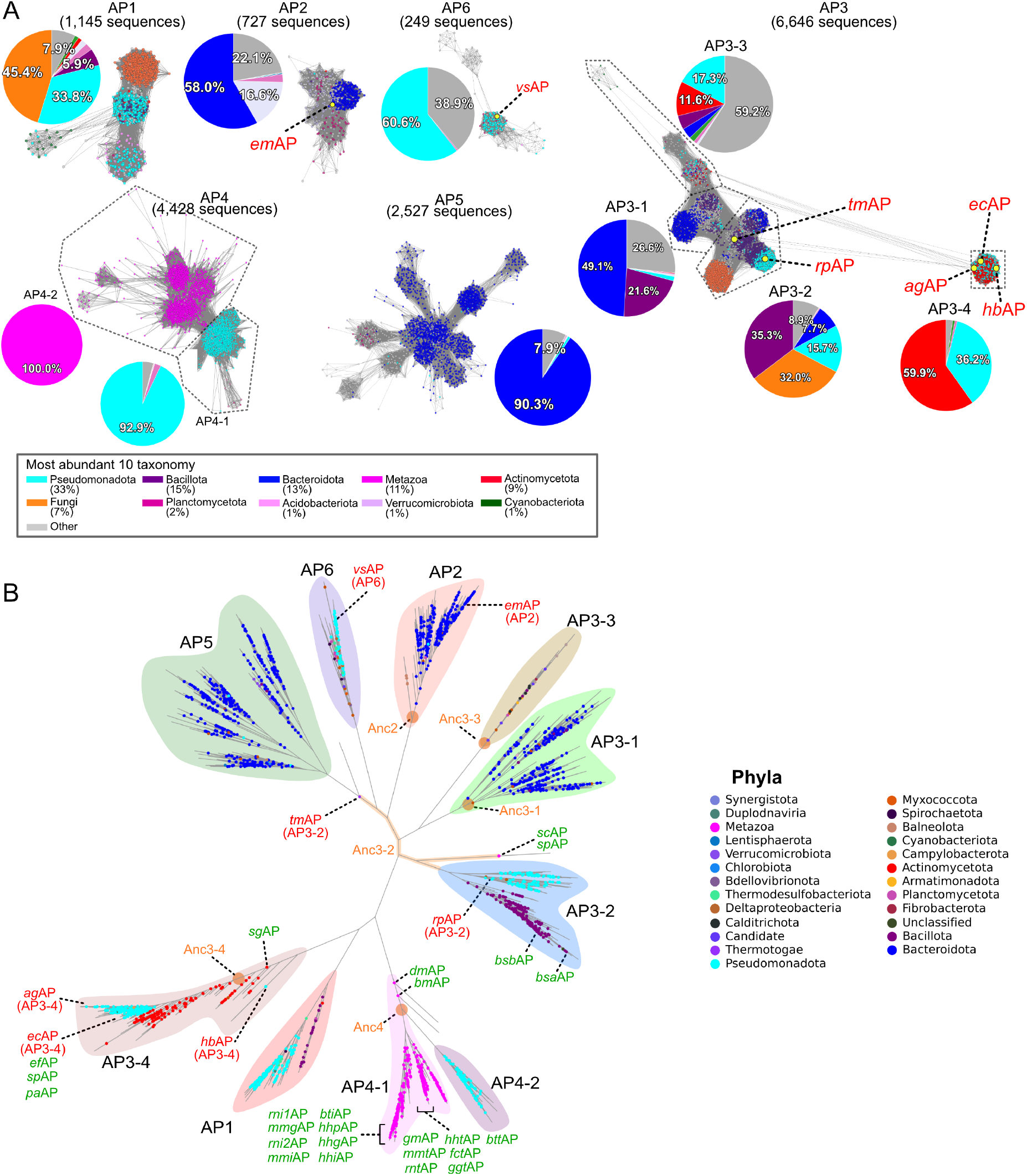
Divergence of AP sequences. (A) SSN analysis of six AP subclusters (AP1-6) identified in the clustering of the AP superfamily database (**Fig. S1A**). The E-value cut-offs for AP1, 2, 5, 6, AP3, and AP4 were 1 × 10^−100^, 1 × 10^−40^, and 1 × 10^−125^, respectively. The ten most abundant phyla are represented by nodes in different colors. AP3 and AP4 are further divided into subgroups (AP3-1, 2, 3, 4 and AP4-1, 2). Pie charts show the phylum distribution within each AP subcluster and subgroup. (B) Phylogenetic tree analysis of AP subcluster. The distribution of the AP subclusters and their associated phyla is mapped onto the phylogenetic tree. Enzymes used for expression and activity measurements, ancestral enzymes, and annotated enzymes are indicated in red, orange, and green, respectively.

Interestingly, the distribution of phyla across the AP subcluster was quite diverse. Especially, *Pseudomonadota, Bacteroidota*, and *Fungi* appeared in multiple AP subclusters: *Pseudomonadota* in AP1, AP3, AP4, and AP6; *Fungi* in AP1 and AP3-2; and *Bacteroidota* in AP2, AP3, and AP5. In some cases, the phylum distribution was resolved at the subgroup level, *e*.*g*., AP4-1 and AP4-2; *Metazoa*/*Pseudomonadota*, AP3-2; *Bacillota/Pseudomonadota/Fungi*, AP3-4; *Actinomycetota*/*Pseudomonadota*. Among 9,588 bacterial genomes in the GTDB database that possess the *phoA* gene, 21% (2,030 genomes) contained multiple copies of *phoA* (**Fig. S7A**), which were clustered in certain bacterial classes such as *Gammaproteobacteria* (in *Pseudomonadota*), and *Bacteroidia* (*in Bacteroidota*) (**Table S1**). We also observed that the multiple copies of the *phoA* gene within these classes were assigned to different AP subclusters (**Fig. S7B**): AP2/AP5, AP3/AP5, AP5/AP5 were commonly found in *Bacteroidota*; AP3/AP4, AP3/AP6, AP3/AP3, and AP4/AP4 were seen in *Pseudomonadota* (**Table S1**). These observations are consistent with our earlier findings that extensive gene gain and loss of the *phoA* gene have occurred among bacteria during their evolutionary divergence (**Fig. S3**).

### Experimental validation of Pt oxidation activity in bacterial AP

A previous study by Yang *et. al*. demonstrated that only *E. coli* alkaline phosphatase (*ec*AP) catalyzed the Pt oxidation reaction, while other APs, including those from *B. subtilis*, calf, and shrimp, did not [24]. However, it remains unclear how prevalent Pt oxidase activity is among other bacterial APs. To address this, we synthesized genes and experimentally characterized thirteen bacterial APs across diverse subgroups (**Fig. S6**). Among these, only seven APs exhibited soluble fractions in SDS-PAGE and measurable phosphomonoesterase (PMEase) activity in lysate assays (**Fig. S8** and **Fig. S9**). Of these seven APs, five belong to AP3 (*Pseudomonadota*: *ec*AP, *Acinetobacter gerneri*: *ag*AP, *Halieaceae bacterium*: *hb*AP, unclassified *Rheinheimera*: *rs*AP, and *Thermotoga maritima*: *tm*AP), one belongs to AP6 (*Pseudomonadota*: unclassified *Vibrio*: *vs*AP), and one belongs to AP2 (*Bacteroidota*: *Elizabethkingia meningoseptica*: *em*AP) (**Fig. 2**). While crystal [35, 36] and predicted structures of these APs revealed several insertion/deletion sites, they all maintained the AP fold with the canonical bimetallic binding site, catalytic residues, and nucleophile serine (**Fig. 3A**). We purified these enzymes and measured both PME and Pt oxidation activities. All the APs exhibited efficient PME activity and, intriguingly, also demonstrated Pt oxidation activity comparable to that of *ec*AP, which provides a growth advantage to the *E. coli* Shuffle strain (Δ*ptxD*, Δ*phoA*) in Pt media (**Fig. 3B-C** and **Fig. S10**).

**Fig. 3.**
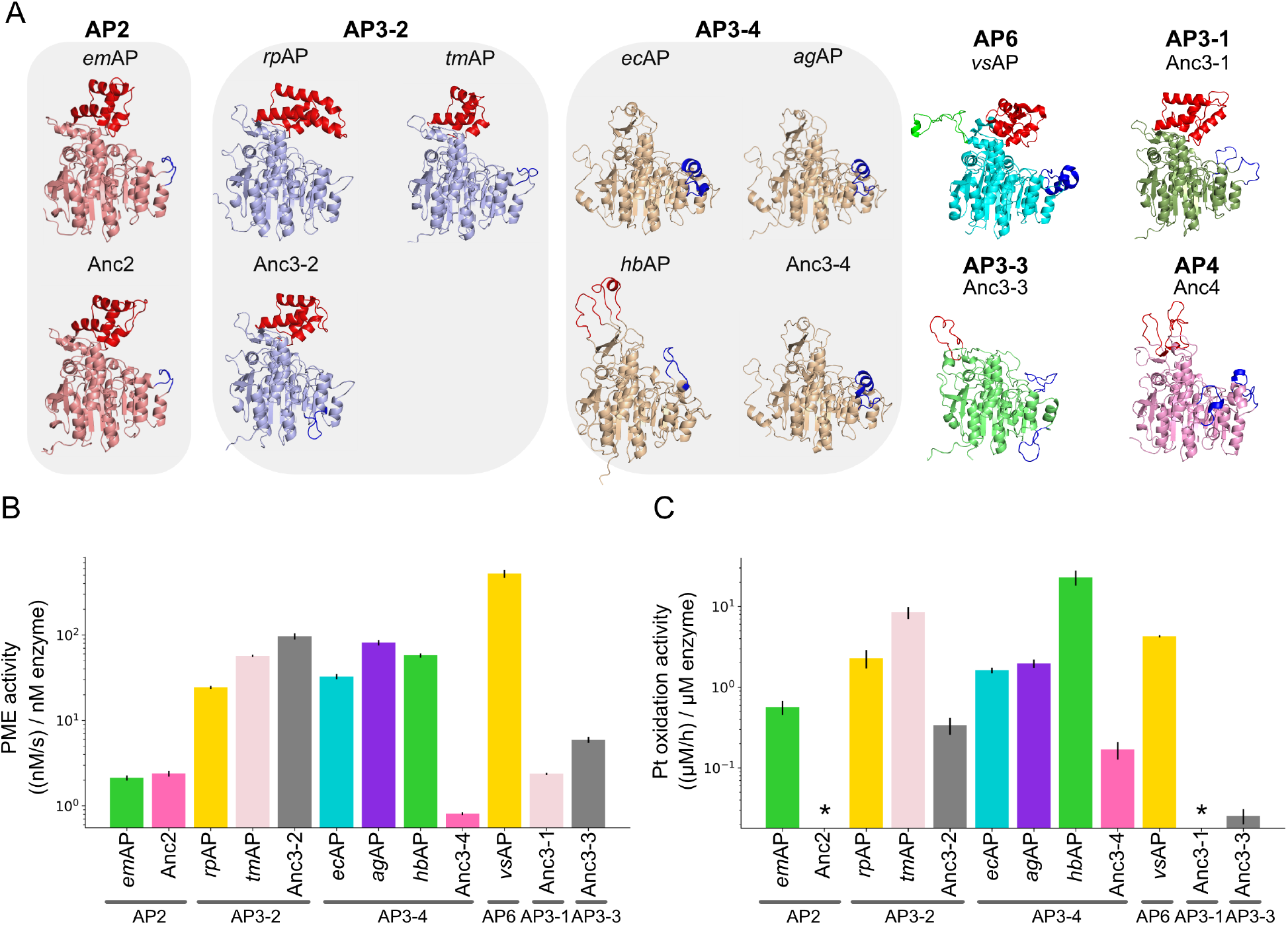
Bulk ensemble kinetic assay of phosphomonoesterase (PMEase) and phosphite (Pt) oxidation functions of homologous APs. (A) Structure of extant and ancestral APs. The *ec*AP (PDB: 1ALK) and *vs*AP (PDB: 3E2D) have crystal structures solved in previous studies. Structures of other APs were estimated using AlphaFold 2. Domains with deletions or insertions are shown in red, blue, and green, respectively. (B), (C) PMEase and Pt oxidation activities of extant and ancestral APs (mean ± *SD*, n = 3). Asterisks indicate enzymes for which activity could not be measured.

We further analyzed the functional profile of inferred ancestral AP enzymes (ancAPs). Using ancestral sequence reconstruction with GRASP [37], we predicted the sequences of common ancestors for several subclusters and subgroups: ancAP2, ancAP3-1, ancAP3-2, ancAP3-3, ancAP3-4, and ancAP4 (**Fig. 2, Fig. 3A**, and **Fig. S11**). All ancAPs, except ancAP4, were expressed in the soluble fraction in *E. coli* (**Fig. S8** and **Fig. S9**). We found that three out of five of the ancAPs—ancAP3-2, ancAP3-3, and ancAP3-4—showed Pt oxidation activity, while all soluble ancAPs exhibited phosphomonoesterase (PMEase) activity (**Fig. 3B-C**). These ancAPs were located at early branching points in the phylogenetic tree (**Fig. 2B**), suggesting that Pt oxidation activity might have been present in the early stages of AP evolution. This function appears to have been inherited in certain lineages and is likely still common among extant APs, although other lineages have lost their ability to oxidize Pt. However, since Pt oxidation by APs is significantly weaker than PMEase activity (**Fig. 3B-C**), it is unlikely to be the primary function of the AP family.

Finally, we analyzed the divergence of Pt oxidation functions among archaeal APs, which are evolutionarily distant from bacterial APs, and PafA, another class of PMEase enzymes belonging to the phosphodiesterase family. We identified 203 *phoA* genes from a total of 4,416 representative archaeal species (**Fig. S12A**), with more than 80% of the sequences containing the canonical bimetallo binding site and nucleophile serine (**Fig. S12B-D**). Compared to bacterial *phoA* genes, archaeal *phoA* genes showed a more limited distribution. Additionally, only one, two, and zero *ptxD, pafA*, and *phnJ* genes, respectively, were identified in archaeal genomes. These findings suggest that archaea may employ different enzymes for PME and Pt oxidation, or, as reported in previous studies, may not utilize Pt as a phosphorus source [21]. We selected two archaeal APs, *Pyrococcus abyssi* [38] and *Halobacterium salinarum* [39], and tested their catalytic activities (**Fig. S12B, E, F**). While *H. salinarum* AP was insoluble in bacterial expression systems, we found that *P. abyssi* AP was soluble (**Fig. S8C**) and exhibited PME activity (0.94 ± 0.07, ((nM/s)/nM enzyme), mean ± *SD*, n = 3, at 60°C), consistent with previous reports [38]. However, we did not observe Pt oxidation activity with purified *P. abyssi* AP, suggesting that Pt oxidation may not be widespread among archaeal APs.

In the case of PafA, we expressed and purified PafA homologs from *Elizabethkingia meningoseptica* (*em*PafA) and *Flavobacterium johnsoniae* (*fj*PafA), with *<u>em</u>*PafA sharing identical metal-binding residues and the same primary function as *ec*AP [26]. While *fj*PafA was insoluble, *em*PafA was expressed in a soluble form (**Fig. S8D**) and exhibited PME activity (315 ± 6 ((nM/s)/nM), mean ± *SD*, n = 3), similar to that of AP. Interestingly, *em*PafA also showed Pt oxidation activity (1.1 ± 0.24 ((nM/h)/nM enzyme, mean ± *SD*, n = 3), suggesting that other enzyme classes with native PME activity may also possess Pt oxidation activity.

### Key active site residues of APs are optimized for Pt oxidation

The Pt oxidation activity by extant and ancestral APs suggests that there may have been selection pressure leading to the retention of this function in the APs. To investigate this further, we analyzed the diversity of active site residues within the AP family. In *ec*AP, six amino acid residues—101, 102, 166, 153, 322, and 328—have been shown to play key roles in PME and promiscuous functions (**Fig. 4A**) [26, 27]. A multiple sequence alignment of the active site residues in each AP subcluster revealed that all APs share the same active site structure, while an overall multiple sequence alignment of APs showed interesting differences between subgroups. While positions 101 (Asp), 102 (Ser), 166 (Arg), and 322 (Glu) are highly conserved across all AP sequences (91-100% conservation), diversities were observed at positions 153 and 328. Position 153 varies between Asp (26%) and His (70%), and position 328 diversifies between His (32%), Lys (14%), and Trp (35%) (**Fig. 4A**). Moreover, AP sequences exhibited specific 153/328 combinations, with three combinations— D153/K328, H153/W328, and H153/H328—found in more than 80% of AP sequences (**Fig. 4B**). Notably, we observed that preference for specific combinations is evident at both the domain and phylum levels, with phyla predominantly possessing one or two specific combinations (**Fig. 4C** and **Fig. S13**). Since the 153/328 combination is conserved at the phylum level, this pattern was also reflected in each subcluster of the SSN, where a specific 153/328 combination is well-preserved (**Fig. 4D** and **Fig. S14**).

**Fig. 4.**
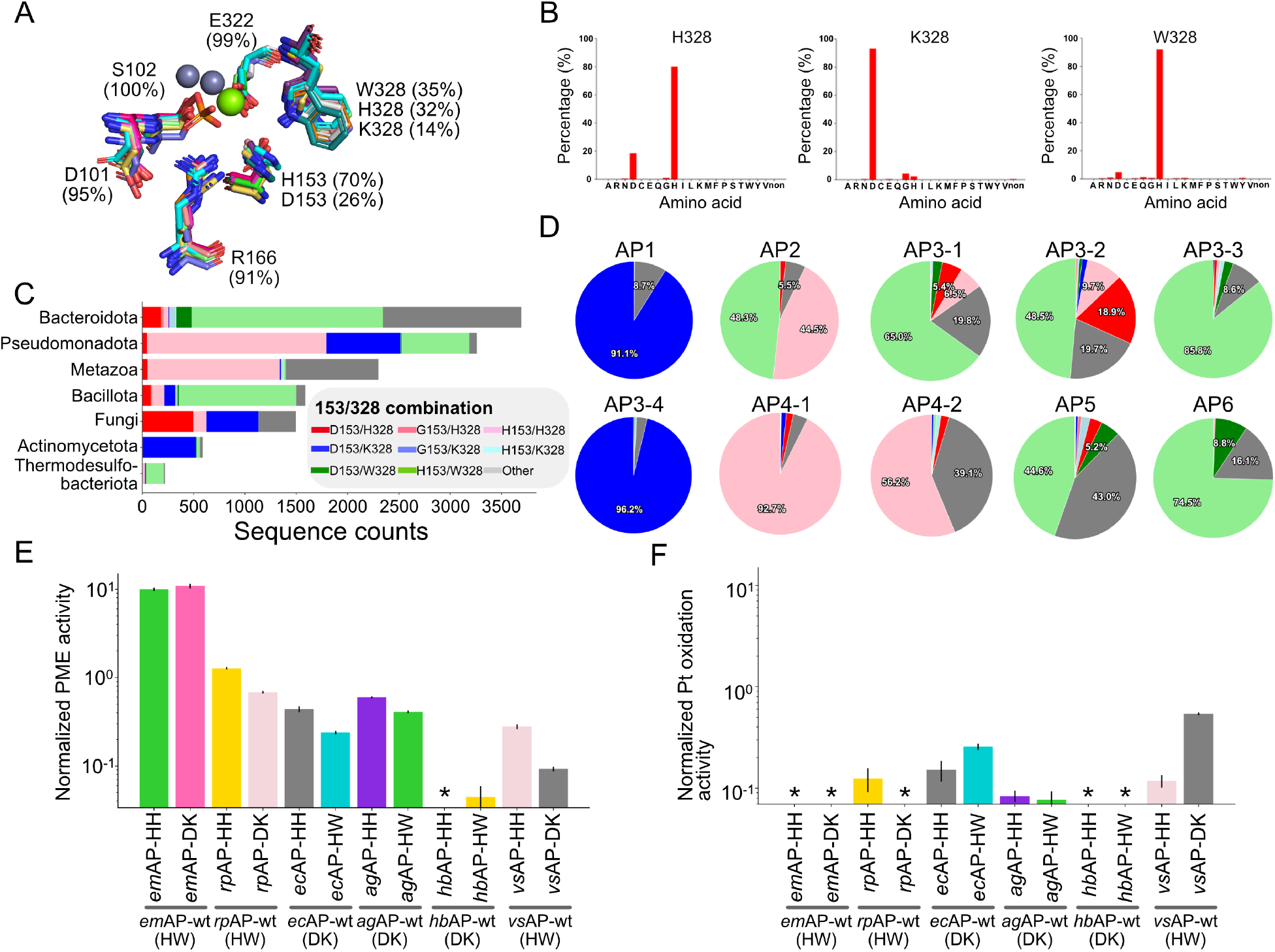
Effect of the 153/328 combination on AP functions. (A) Alignment of the active site structures. Extant and ancestral enzymes shown in **Fig. 3A** were used for the alignment. Dark and green spheres represent zinc and magnesium ions identified by the crystal structure of *ec*AP (PDB: 1ALK). (B) Ratio of the specific 153/328 combination in the AP subclusters. Each graph shows the percentage of the amino acid residue at position 153 when the residue at position 328 is H, K, or W, respectively. (C) Sequence count of the 153/328 combination in the most abundant seven phyla. (D) Ratio of the specific 153/328 combination in each AP subcluster. The color code is the same as in **Fig. 4C**. (E), (F) PMEase and Pt oxidation activities of swapping mutants (mean ± *SD*, n = 3). Activities of the mutants were normalized to those of their wild-type enzymes. Asterisks indicate enzymes for which activity could hardly be measured.

Is the conservation of the 153/328 combination a consequence of functional constraints? In other words, are specific combinations in each subcluster necessary for the function of APs? To analyze the mutational effects of the 153/328 combinations, we generated a series of swapping mutants based on the six APs and measured their PMEase and Pt oxidation activities (**Fig. 4E-F**). The wild-type sequences of *em*AP, *rp*AP, and *vs*AP contain H153/W328, while those of *ec*AP, *ag*AP, and *hb*AP have D153/K328. We selected the most dominant swapping mutations—H153/H328, H153/W328, and D153/K328. Swapping combinations strongly affected PMEase activity in various ways depending on the genetic background. For example, the swapping mutations negatively affected the PMEase activities of *ec*AP, *vs*AP, and *hb*AP, while they positively influenced *em*AP. *rp*AP exhibited both positive and negative effects depending on the mutations (**Fig. 4E**). Mutations at positions 153 and 328 resulted in various effects, including changes in the rate-limiting step, altered magnesium affinity, and shifts in phosphate positioning [27, 40]. The complex interplay of these effects, along with interactions with different surrounding residues in homologous enzymes, leads to enzyme-specific variations in catalytic properties. In contrast, swapping mutations consistently showed negative effects on Pt oxidation activity across all tested APs (**Fig. 4F**). For *ec*AP, *ag*AP, and *rp*AP, Pt oxidation decreased by more than fivefold. Moreover, Pt oxidation could not be detected in the mutated *hb*AP and *em*AP, even though these mutations had a positive impact on the PMEase activity of *em*AP. Thus, the wild-type combination in each enzyme appears to be somewhat subjected to selection pressure and optimized for Pt oxidation, but it is not essential for PMEase activity. Since the balance of activities was easily disrupted by mutations in the active site, organisms may frequently use AP that already retain an optimized functional spectrum for organism growth, as reflected by their frequent transfer (**Fig. 2**).

## Discussion

This study highlights the overlooked ecological significance of the promiscuous Pt oxidation function and its potential contribution to the global phosphorus cycle. While microbes have evolved to utilize diverse phosphorus compounds, early life likely first relied on the most readily available forms—primarily inorganic phosphates—to expand their metabolic capabilities. However, the limited solubility of phosphate and the intense competition for its assimilation may have driven the evolution of diverse enzymatic pathways to process various DOP sources [1-3]. Given that nucleic acids and numerous phosphorylated metabolites are indispensable for organism growth [41, 42], phosphatases would have emerged in early life to recycle and produce bioavailable phosphate. Indeed, a previous study suggested that inferred ancestral metabolic pathways in LUCA contained several phosphatases, including AP [32]. Our observations suggest that these phosphatases may have provided ancient microbes with the ability to access other phosphorus sources, *i*.*e*., Pt, via their multifunctionality [25-27, 42]. Despite the low levels of Pt oxidase activity observed in APs, they were sufficient levels to support *E. coli* growth in Pt-containing media (**Fig. S10**) [24], suggesting potential physiological roles in natural environments, particularly where microbes have access to a mixture of diverse phosphorus sources, including inorganic phosphate, DOPs, and Pt [1-3, 5, 7]. A more efficient and specialized Pt oxidase, PtxD, appeared later than AP and likely spread among bacteria that require much higher catalytic Pt oxidase activity [1, 18-20].

Pt concentration gradually declined as Earth’s atmosphere oxidized after the GOE, with reduced inputs, such as meteorite impacts and lightning events, contributing to this decrease [1, 3, 12, 43, 44]. However, Pt continues to be produced abiotically [3, 45-48] and remains ubiquitous, including in anoxic environments [18-20, 49]. Thus, it is highly likely that accessing phosphite as a phosphorus source can give an advantage to diverse modern bacteria. In this study, we found that 47% of bacterial genomes (15,305 out of 32,255) harbor at least one gene encoding a potential Pt oxidase, including the *phoA, pafA, ptxD*, or *phnJ* genes (**Fig. 1B**). Additionally, given the structural similarity of their active sites [25, 26], it is also possible that other phosphatases can catalyze Pt oxidation, which may explain previous observations that 10–67% of bacterial isolates from environmental samples are capable of growing in Pt-containing media [22]. Further characterization of promiscuous Pt oxidation by phosphatases and CP lyases, along with investigation of the distribution of environmental microbes encoding these genes, is expected to provide new insights into the role of Pt in bacterial phosphorus metabolism.

## Acknowledgment

The authors are grateful to Dan Kehila, Lidia Piatnitca, and Santanu Sasidharan for technical assistance and to Liam M. Longo and members of the Tokuriki lab for helpful discussions and comments on the manuscript. M.S. is supported by the Japan Society for the Promotion of Science (JSPS) Overseas Research Fellowship. N.K. received the support of the Grant-in-Aid for JSPS Fellows from JSPS (22J20318). J.Z.C. received the support of the Human Frontier Science Program postdoctoral fellowship (LT0048/2023-L). N.T. is supported by Natural Sciences and Engineering Research Council of Canada (NSERC) / Discovery Grants Program (RGPIN-2023-05135) and the Human Frontier Science Program (HFSP) Research Grant (RGP0054/2020).

## Author contributions

M.S. conceived the project, designed experiments, performed experiments, analyzed results, and wrote the manuscript; N.K., S.G., and J.Z.C. performed experiments, analyzed results, and wrote the manuscript; N.T. conceived the project, designed experiments and wrote the manuscript.

## References

1. Figueroa IA, Coates JD: Microbial Phosphite Oxidation and Its Potential Role in the Global Phosphorus and Carbon Cycles. Adv Appl Microbiol 2017, 98:93–117.

2. Nicholls JWF, Chin JP, Williams TA, Lenton TM, O’Flaherty V, McGrath JW: On the potential roles of phosphorus in the early evolution of energy metabolism. Front Microbiol 2023, 14:1239189.

3. Duhamel S: The microbial phosphorus cycle in aquatic ecosystems. Nat Rev Microbiol 2024, 23:239–255.

4. Margalef O, Sardans J, Fernández-Martínez M, Molowny-Horas R, Janssens IA, Ciais P, Goll D, Richter A, Obersteiner M, Asensio D, et al.: Global patterns of phosphatase activity in natural soils. Sci Rep 2017, 7:1337.

5. Srivastava A, Saavedra DEM, Thomson B, García JAL, Zhao Z, Patrick WM, Herndl GJ, Baltar F: Enzyme promiscuity in natural environments: alkaline phosphatase in the ocean. ISME J 2021, 15:3375–3383.

6. Lidbury IDEA, Scanlan DJ, Murphy ARJ, Christie-Oleza JA, Aguilo-Ferretjans MM, Hitchcock A, Daniell TJ: A widely distributed phosphate-insensitive phosphatase presents a route for rapid organophosphorus remineralization in the biosphere. Proc Natl Acad Sci U S A 2022, 119:e2118122119.

7. Despotovic D, Aharon E, Trofimyuk O, Dubovetskyi A, Cherukuri KP, Ashani Y, Eliason O, Sperfeld M, Leader H, Castelli A, et al.: Utilization of diverse organophosphorus pollutants by marine bacteria. Proc Natl Acad Sci U S A 2022, 119:e2203604119.

8. White AK, Metcalf WW: Microbial metabolism of reduced phosphorus compounds. Annu Rev Microbiol 2007, 61:379–400.

9. Sosa OA, Repeta DJ, DeLong EF, Ashkezari MD, Karl DM: Phosphate-limited ocean regions select for bacterial populations enriched in the carbon–phosphorus lyase pathway for phosphonate degradation. Environ Microbiol 2019, 21:2402–2414.

10. Tapia-Torres Y, Olmedo-Álvarez G: Life on Phosphite: A Metagenomics Tale. Trends Microbiol 2018, 26:170–172.

11. Schwartz AW: Phosphorus in prebiotic chemistry. Philos Trans R Soc B Biol Sci 2006, 361:1743–1749.

12. Pasek MA: Rethinking early Earth phosphorus geochemistry. Proc Natl Acad Sci U S A 2008, 105:853–858.

13. Pasek M, Block K: Lightning-induced reduction of phosphorus oxidation state. Nat Geosci 2009, 2:553–556.

14. Pech H, Henry A, Khachikian CS, Salmassi TM, Hanrahan G, Foster KL: Detection of geothermal phosphite using high-performance liquid chromatography. Environ Sci Technol 2009, 43:7671–7675.

15. Garcia Costas AM, White AK, Metcalf WW: Purification and Characterization of a Novel Phosphorus-oxidizing Enzyme from Pseudomonas stutzeri WM88. J Biol Chem 2001, 276:17429–17436.

16. Zou Y, Zhang H, Brunzelle JS, Johannes TW, Woodyer R, Hung JE, Nair N, Van Der Donk WA, Zhao H, Nair SK: Crystal structures of phosphite dehydrogenase provide insights into nicotinamide cofactor regeneration. Biochemistry 2012, 51:4263–4270.

17. Metcalf WW, Wolfe RS: Molecular genetic analysis of phosphite and hypophosphite oxidation by Pseudomonas stutzeri WM88. J Bacteriol 1998, 180:5547–5558.

18. Poehlein A, Daniel R, Schink B, Simeonova DD: Life based on phosphite: A genome-guided analysis of Desulfotignum phosphitoxidans. BMC Genomics 2013, 14:753.

19. Figueroa IA, Barnum TP, Somasekhar PY, Carlström CI, Engelbrektson AL, Coates JD: Metagenomics-guided analysis of microbial chemolithoautotrophic phosphite oxidation yields evidence of a seventh natural CO2 fixation pathway. Proc Natl Acad Sci U S A 2018, 115:E92–E101.

20. Ewens SD, Gomberg AFS, Barnum TP, Borton MA, Carlson HK, Wrighton KC, Coates JD: The diversity and evolution of microbial dissimilatory phosphite oxidation. Proc Natl Acad Sci U S A 2021, 118:e2020024118.

21. Boden JS, Zhong J, Anderson RE, Stüeken EE: Timing the evolution of phosphorus-cycling enzymes through geological time using phylogenomics. Nat Commun 2024, 15:3703.

22. Stone BL, White AK: Most probable number quantification of hypophosphite and phosphite oxidizing bacteria in natural aquatic and terrestrial environments. Arch Microbiol 2012, 194:223–228.

23. Metcalf WW, Wanner BL: Involvement of the Escherichia coli phn (psiD) gene cluster in assimilation of phosphorus in the form of phosphonates, phosphite, P(i) esters, and P(i). J Bacteriol 1991, 173:587–600.

24. Yang K, Metcalf WW: A new activity for an old enzyme: Escherichia coli bacterial alkaline phosphatase is a phosphite-dependent hydrogenase. Proc Natl Acad Sci U S A 2004, 101:7919–7924.

25. Mohamed MF, Hollfelder F: Efficient, crosswise catalytic promiscuity among enzymes that catalyze phosphoryl transfer. Biochim Biophys Acta - Proteins Proteomics 2013, 1834:417–424.

26. Sunden F, Alsadhan I, Lyubimov A, Doukov T, Swan J, Herschlag D: Differential catalytic promiscuity of the alkaline phosphatase superfamily bimetallo core reveals mechanistic features underlying enzyme evolution. J Biol Chem 2017, 292:20960–20974.

27. Sakuma M, Honda S, Ueno H, Tabata K V., Miyazaki K, Tokuriki N, Noji H: Genetic Perturbation Alters Functional Substates in Alkaline Phosphatase. J Am Chem Soc 2023, 145:2806–2814.

28. White AK, Metcalf WW: Two C-P lyase operons in Pseudomonas stutzeri and their roles in the oxidation of phosphonates, phosphite, and hypophosphite. J Bacteriol 2004, 186:4730–4739.

29. Konno N, Iwasaki W: Machine learning enables prediction of metabolic system evolution in bacteria. Sci Adv 2023, 9:eadc9130.

30. Gholipour S, Chen J, Lee D, Tokuriki N: The global β-lactam resistome revealed by comprehensive sequence analysis. bioRxiv 2024, 10.1101/2024.03.01.583042

31. Van Dien SJ, Keasling JD: A dynamic model of the Escherichia coli phosphate-starvation response. J Theor Biol 1998, 190:37–49.

32. Moody ERR, Álvarez-Carretero S, Mahendrarajah TA, Clark JW, Betts HC, Dombrowski N, Szánthó LL, Boyle RA, Daines S, Chen X, et al.: The nature of the last universal common ancestor and its impact on the early Earth system. Nat Ecol Evol 2024, 8:1654–1666.

33. Liu C, Kenney T, Beiko RG, Gu H: The Community Coevolution Model with Application to the Study of Evolutionary Relationships between Genes Based on Phylogenetic Profiles. Syst Biol 2023, 72:559–574.

34. Ishikawa SA, Zhukova A, Iwasaki W, Gascuel O, Pupko T: A Fast Likelihood Method to Reconstruct and Visualize Ancestral Scenarios. Mol Biol Evol 2019, 36:2069–2085.

35. Helland R, Larsen RL, Ásgeirsson B: The 1.4 Å crystal structure of the large and cold-active Vibrio sp. alkaline phosphatase. Biochim Biophys Acta - Proteins Proteomics 2009, 1794:297–308.

36. Kim EE, Wyckoff HW: Reaction Mechanism of Alkaline Phosphatase Based on Crystal Structures. J Mol Biol 1991, 218:449–464.

37. Foley G, Mora A, Ross CM, Bottoms S, Sützl L, Lamprecht ML, Zaugg J, Essebier A, Balderson B, Newell R, et al.: Engineering indel and substitution variants of diverse and ancient enzymes using Graphical Representation of Ancestral Sequence Predictions (GRASP). PLoS Comput Biol 2022, 18:e1010633.

38. Zappa S, Rolland JL, Flament D, Gueguen Y, Boudrant J, Dietrich J: Characterization of a Highly Thermostable Alkaline Phosphatase from the Euryarchaeon Pyrococcus abyssi. Appl Environ Microbiol 2001, 67:4504–4511.

39. Wende A, Johansson P, Vollrath R, Dyall-Smith M, Oesterhelt D, Grininger M: Structural and biochemical characterization of a halophilic archaeal alkaline phosphatase. J Mol Biol 2010, 400:52–62.

40. Holtz KM, Kantrowitz ER: The mechanism of the alkaline phosphatase reaction: Insights from NMR, crystallography and site-specific mutagenesis. FEBS Lett 1999, 462:7–11.

41. Xavier JC, Gerhards RE, Wimmer JLE, Brueckner J, Tria FDK, Martin WF: The metabolic network of the last bacterial common ancestor. Commun Biol 2021, 4:413.

42. Kuznetsova E, Proudfoot M, Gonzalez CF, Brown G, Omelchenko M V., Borozan I, Carmel L, Wolf YI, Mori H, Savchenko A V., et al.: Genome-wide analysis of substrate specificities of the Escherichia coli haloacid dehalogenase-like phosphatase family. J Biol Chem 2006, 281:36149–36161.

43. Pasek MA, Kee TP: Origins of Life: The Primal Self-Organization. (Springer, 2011). 10.1007/978-3-642-21625-1.

44. Pasek MA, Sampson JM, Atlas Z: Redox chemistry in the phosphorus biogeochemical cycle. Proc Natl Acad Sci U S A 2014, 111:15468–15473.

45. Freeman S, Irwin WJ, Schwalbe CH: Synthesis and Hydrolysis Studies of Phosphonopyruvate. J. Chem. Soc. Perkin Trans 1991, 2:263–267.

46. Momcheva I, Brammer GB, Fynbo JPU, Kashikawa N, Dey A, Matsuda Y, Zabludoff A, Tremonti C, Eisenstein D, Overzier R, et al.: Major role of planktonic phosphate reduction in the marine phosphorus redox cycle. Science 2015, 348:783–785.

47. Bisson C, Adams NBP, Stevenson B, Brindley AA, Polyviou D, Bibby TS, Baker PJ, Hunter CN, Hitchcock A: The molecular basis of phosphite and hypophosphite recognition by ABC-transporters. Nat Commun 2017, 8:1746.

48. Pasek M: A role for phosphorus redox in emerging and modern biochemistry. Curr Opin Chem Biol 2019, 49:53–58.

49. Liu W, Zhang Y, Yu M, Xu J, Du H, Zhang R, Wu D, Xie X: Role of phosphite in the environmental phosphorus cycle. Sci Total Environ 2023, 881:163463.

