## Supplementary information for "Widespread promiscuous alkaline phosphatases underscore early microbial phosphite utilization"

**Supplementary Information content:**

**Figure S1** Sequence clustering analysis of AP superfamily, PtxD, PhnJ, and PstS.

**Figure S2** Analysis of co-occurrence and mutual exclusivity among enzyme genes

**Figure S3** Ancestral state estimation in the phylogenetic tree of bacterial genomes

**Figure S4** Amino acid residue ratio of the bimetallo site and nucleophilic serine in each of the AP subclusters

**Figure S5** Catalytic amino acid residue ratio in each of the AP subclusters

**Figure S6** Distribution of annotated AP sequences on SSN

**Figure S7** Distribution of multiple copies of *phoA* genes and AP subclusters on the phylogenetic tree of bacterial genomes

**Figure S8** SDS-PAGE gels of expressed AP and PafA

**Figure S9** Lysate PMEase activity of soluble APs

**Figure S10** Bacterial growth assay of T7 Shuffle bacteria (Δ*ptxD*, Δ*phoA*) with *ec*AP expressing plasmid and plasmid-free control

**Figure S11** Ancestral enzyme reconstruction in each of the AP subclusters.

**Figure S12** Phylogenetic tree and SSN analysis of Archaeal APs.

**Figure S13** Selection of the combination of positions 153 and 328 in each domain and phylum within the AP subcluster

**Figure S14** Mapping positions 153 and 328 combinations on SSN in the AP subclusters

**Table S1** Sequences used in this study and data on multiple copies of the *phoA* gene

**Materials and methods**

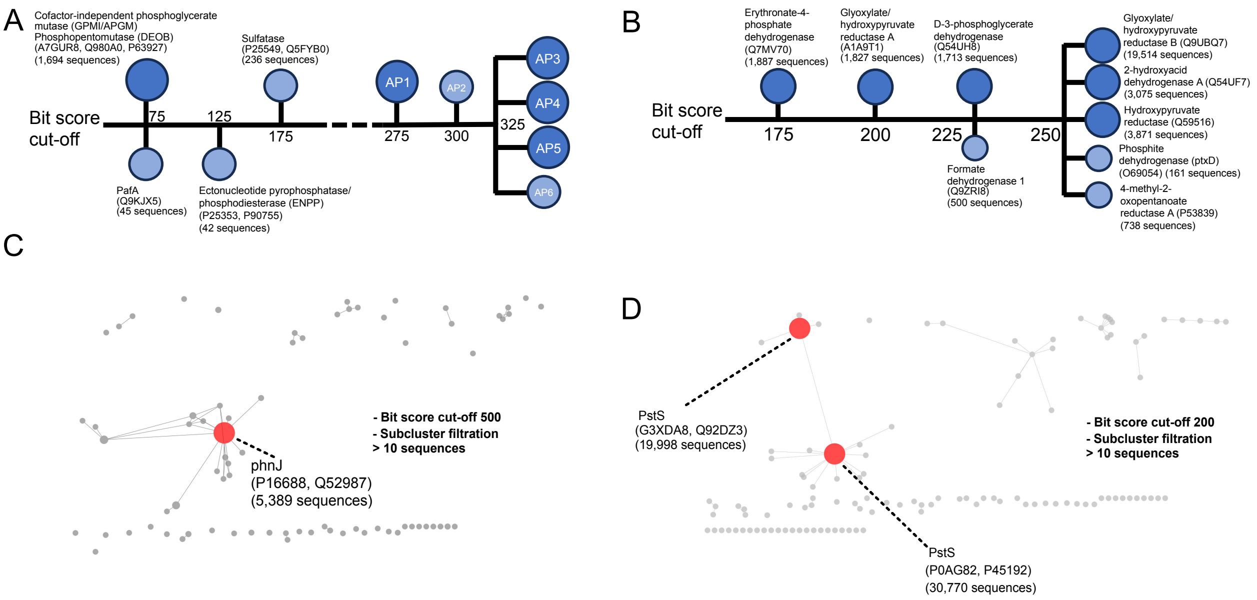


**Figure S1** Sequence clustering analysis of AP superfamily (A), PtxD (B), PhnJ (C), and PstS (D). Clusters were identified based on reviewed sequences from Swiss-Prot in UniProt.

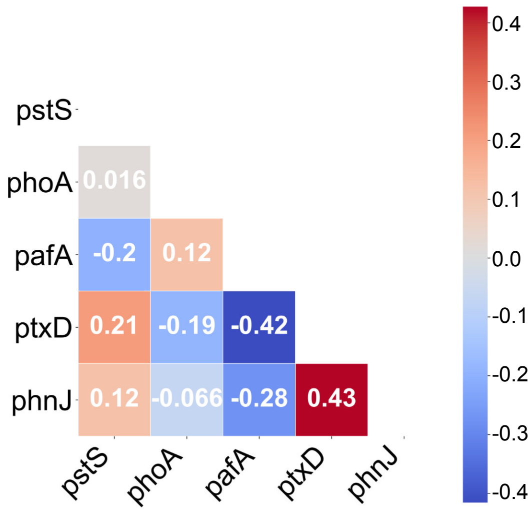


**Figure S2** Analysis of co-occurrence and mutual exclusivity among enzyme genes. EvolCCM was employed to analyze all gene combinations among the *pstS*, *phoA*, *pafA*, *ptxD*, and *phnJ* genes. All gene pairs except the *phoA* and *pstS* (p-value >0.1) and *phoA* and *phnJ* genes (p-value <0.05) showed p-values less than 0.001.

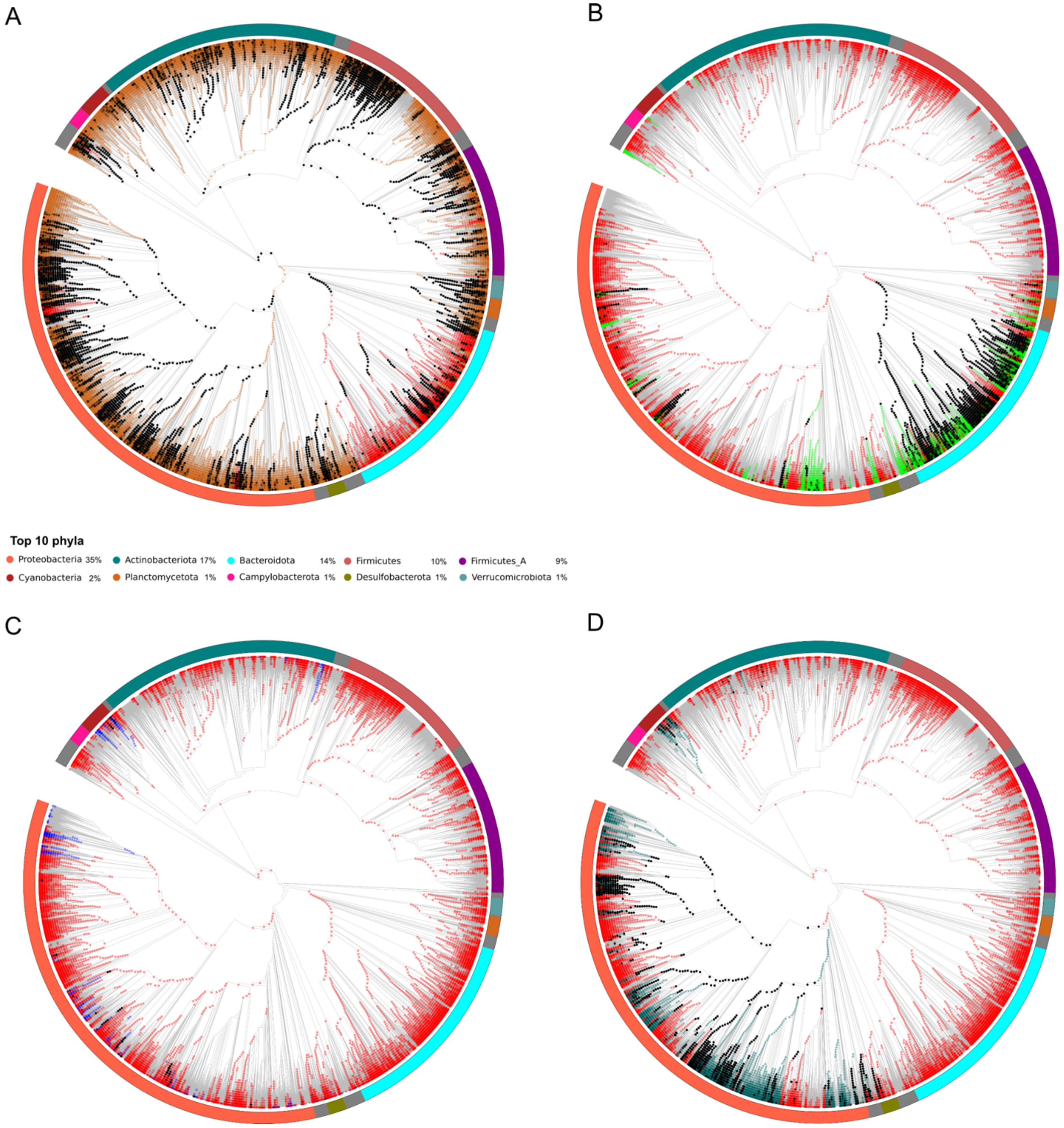


**Figure S3** Ancestral state estimation in the phylogenetic tree of the bacterial genomes. Mapping ancestral state of the *phoA* (red circles) and *pstS* (brown) (A), *pafA* (green) (B), *ptxD* (blue) (C), and *phnJ* (dark green) (D) genes estimated by using PastML. Black-colored nodes indicate that both enzymes have ancestral states.

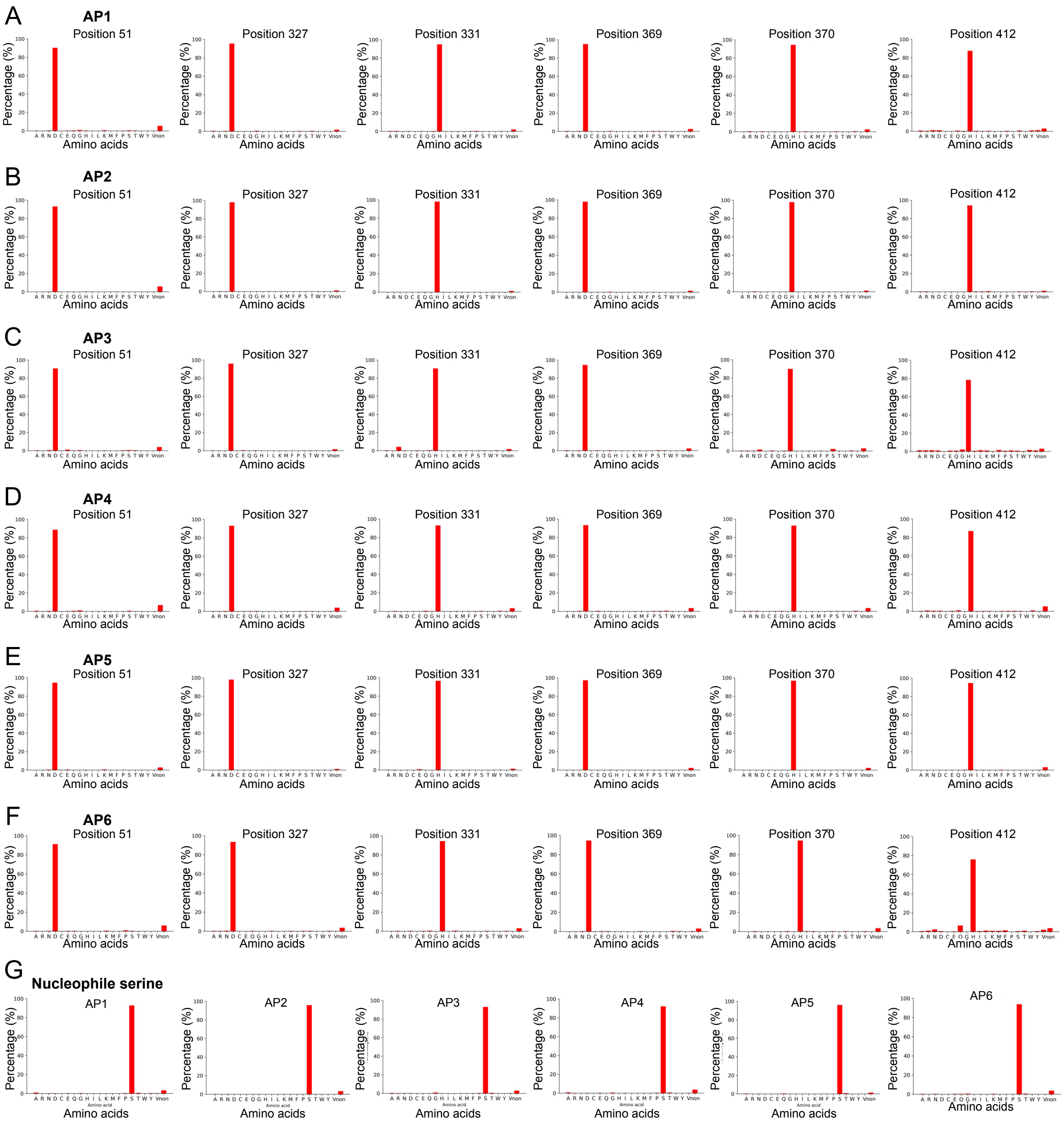


**Figure S4** Amino acid residue ratio of the bimetallo site and nucleophilic serine in each of the AP subclusters. Those positions (51, 327, 331, 369, 370, and 412 in *ec*AP) were $Zn^{2+}$-bimetallo sites. Nucleophile serine was observed at position 102 in *ec*AP.

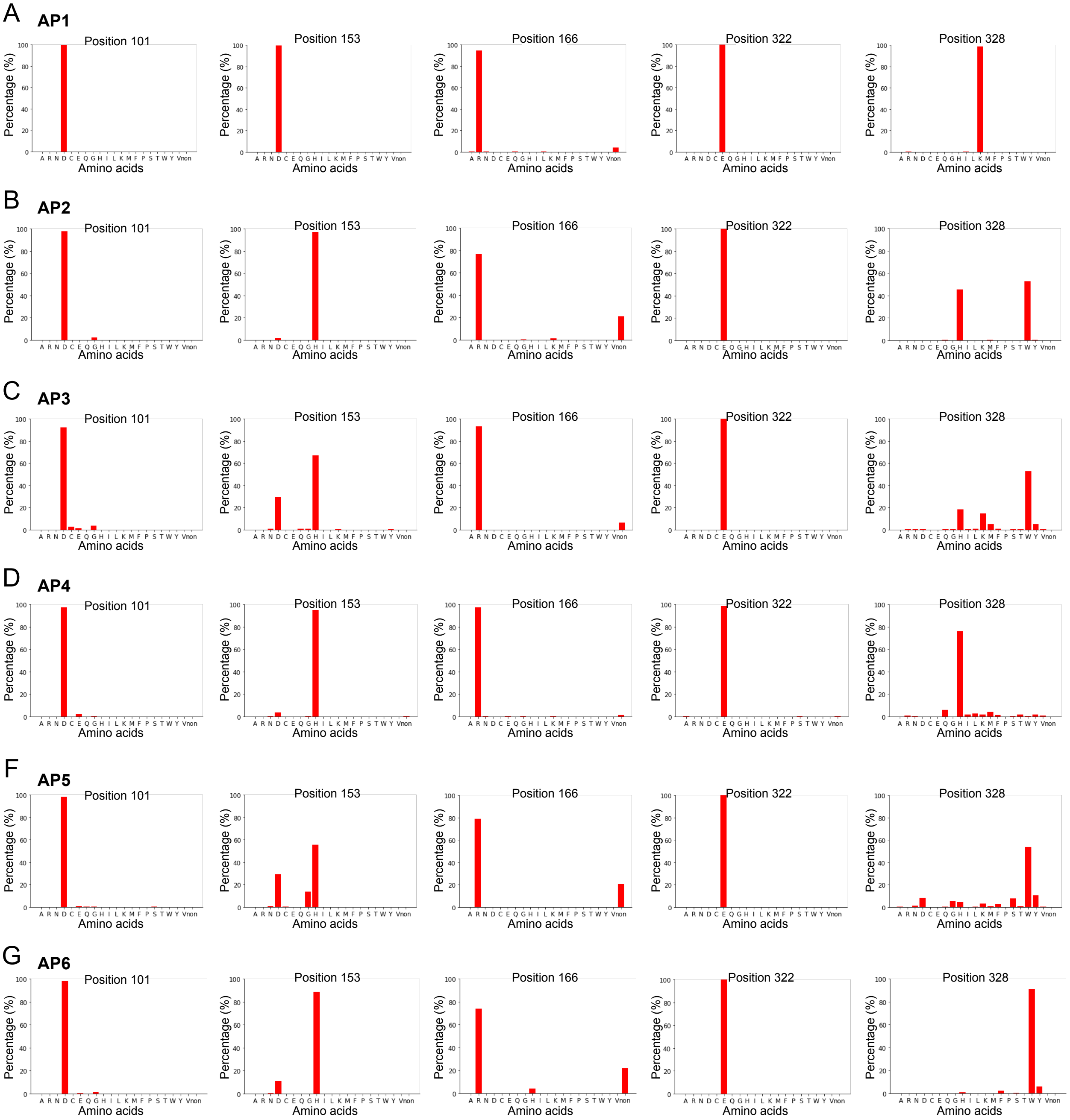


**Figure S5** Catalytic amino acid residue ratio in each of the AP subclusters. The positions of each amino acid are shown in **Fig. 4A**.

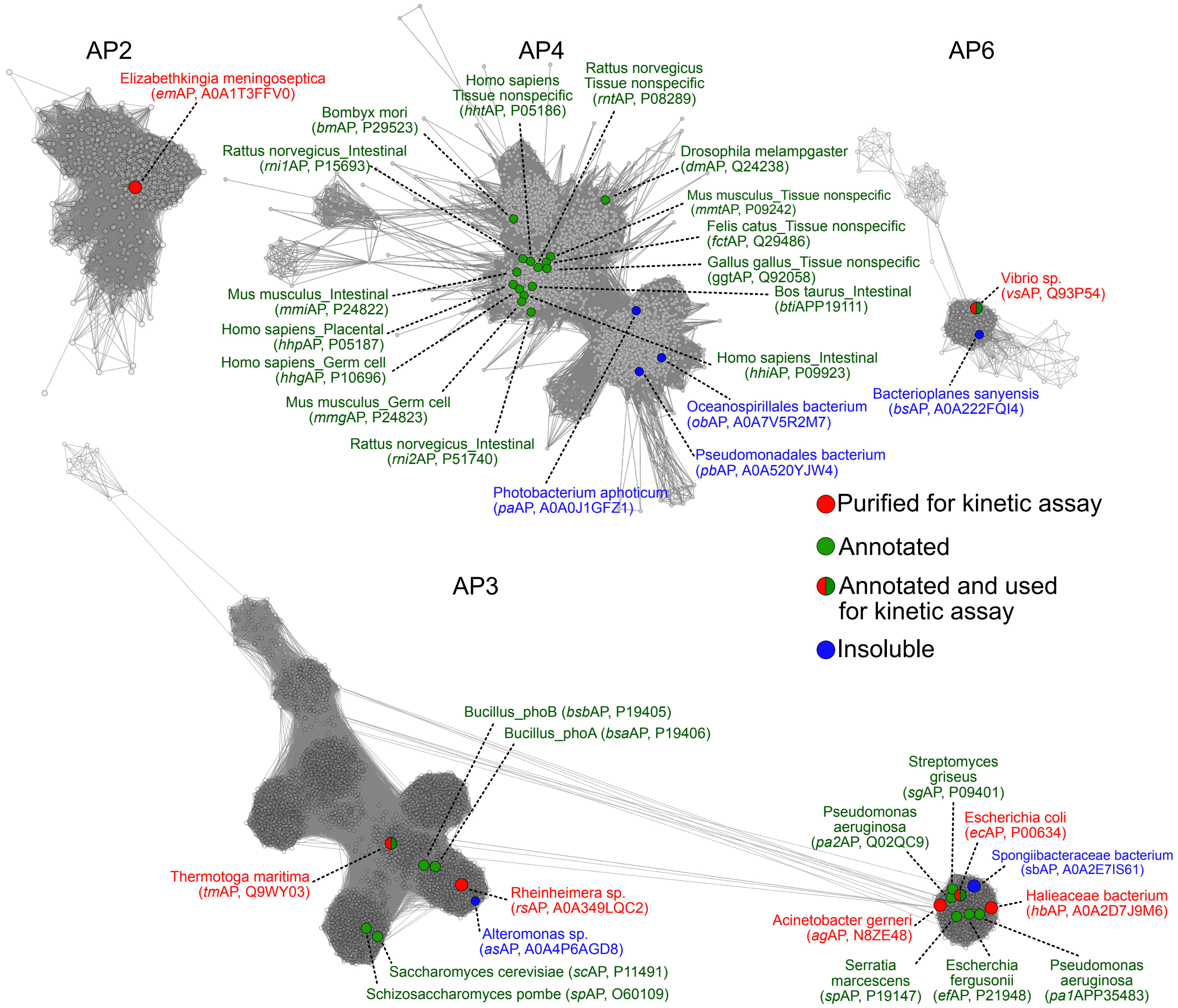


**Figure S6** Distribution of annotated AP sequences on SSN. The green, red, blue, and half green and red nodes indicate APs with annotations, sequences without annotations but used for kinetic assay, insoluble APs without annotation, and sequences with annotations and used for kinetic assay, respectively.

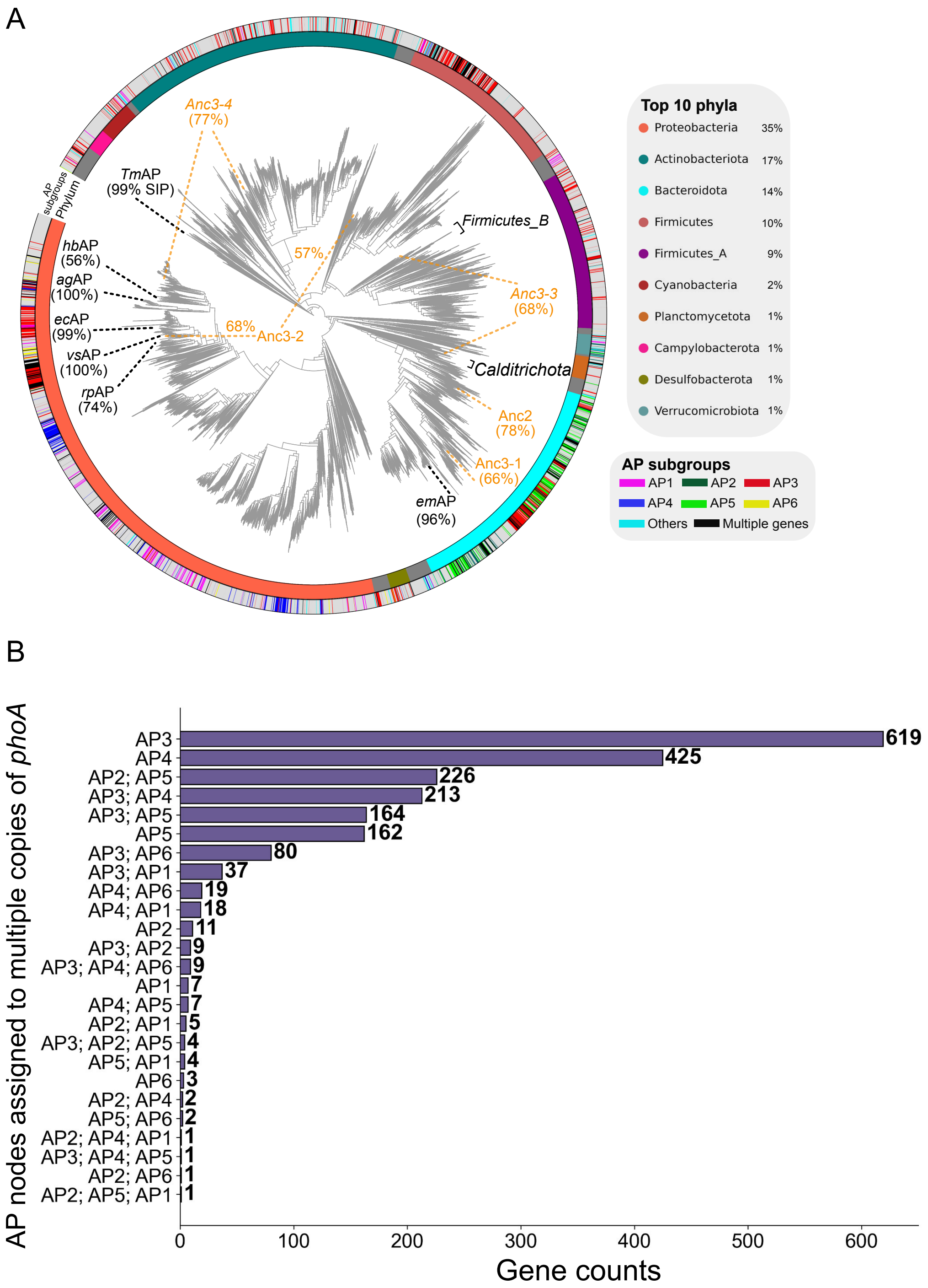


**Figure S7** Distribution of multiple copies of the *phoA* gene and AP subclusters on the phylogenetic tree of bacterial genomes. (A) Clustering the *phoA* gene in GTDB sequences with the AP database identified which subclusters they belong to. The phylogenetic tree also shows the positions of the twelve APs exhibiting Pt oxidation activity along with the sequences that have the highest sequence identity percentage (SIP) to the same as **Fig. 1B**. (B) The gene counts of AP nodes assigned to multiple copies of the *phoA* gene.

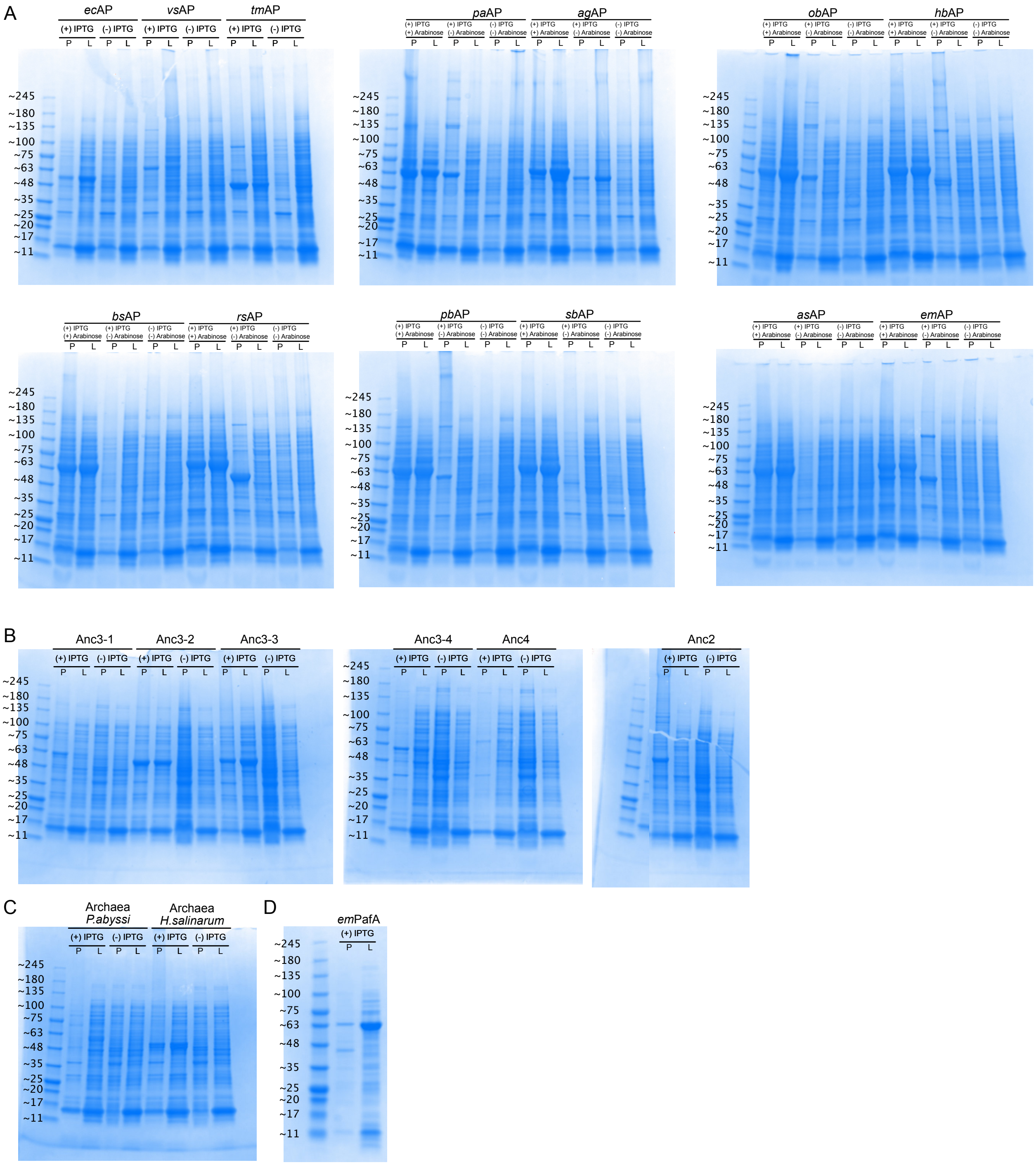


**Figure S8** SDS-PAGE gels of expressed APs and PafA. (A)-(D) The SDS-PAGE gels of extant APs, ancestral APs, Archaeal APs, and PafA, respectively. P and L indicate pellet and lysate. IPTG and Arabinose were used for expressing AP or co-expressing AP and GroEL/ES chaperonin, respectively.

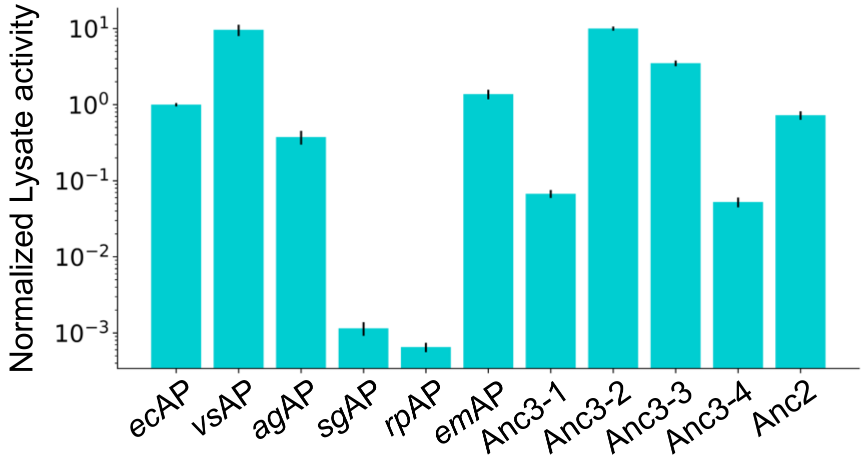


**Figure S9** Lysate PMEase activity of soluble APs. Lysate activity was normalized with that of *ec*AP.

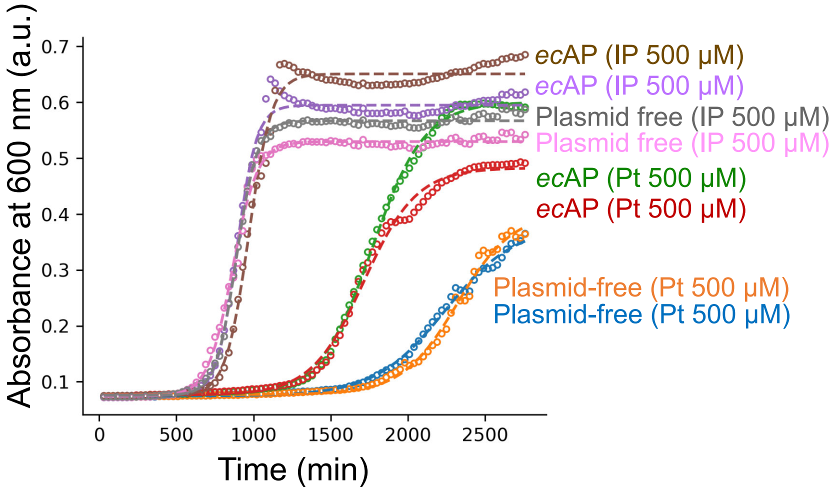


**Figure S10** Bacterial growth assay of T7 Shuffle bacteria (Δ*ptxD*, Δ*phoA*) with *ec*AP expressing plasmid and plasmid-free control. Bacterial growth was measured in media containing 500 μM inorganic phosphate (IP) or phosphite (Pt). Dashed lines represent logistic curve fitting used to estimate the growth rate (r) and the time to reach 50% of maximum growth ($x_0$). The calculated parameters (r, $x_0$) were as follows (mean ± *SD*, n = 2): *ec*AP (IP), r = 0.015 ± 0.002 and $x_0$ = 930 ± 56; *ec*AP (Pt), r = 0.0060 ± 0.00002 and $x_0$ = 1,741 ± 37; plasmid-free (IP), r = 0.014 ± 0.002 and $x_0$ = 876 ± 18; plasmid-free (Pt), r = 0.0050 ± 0.0008 and $x_0$ = 2,281 ± 56.

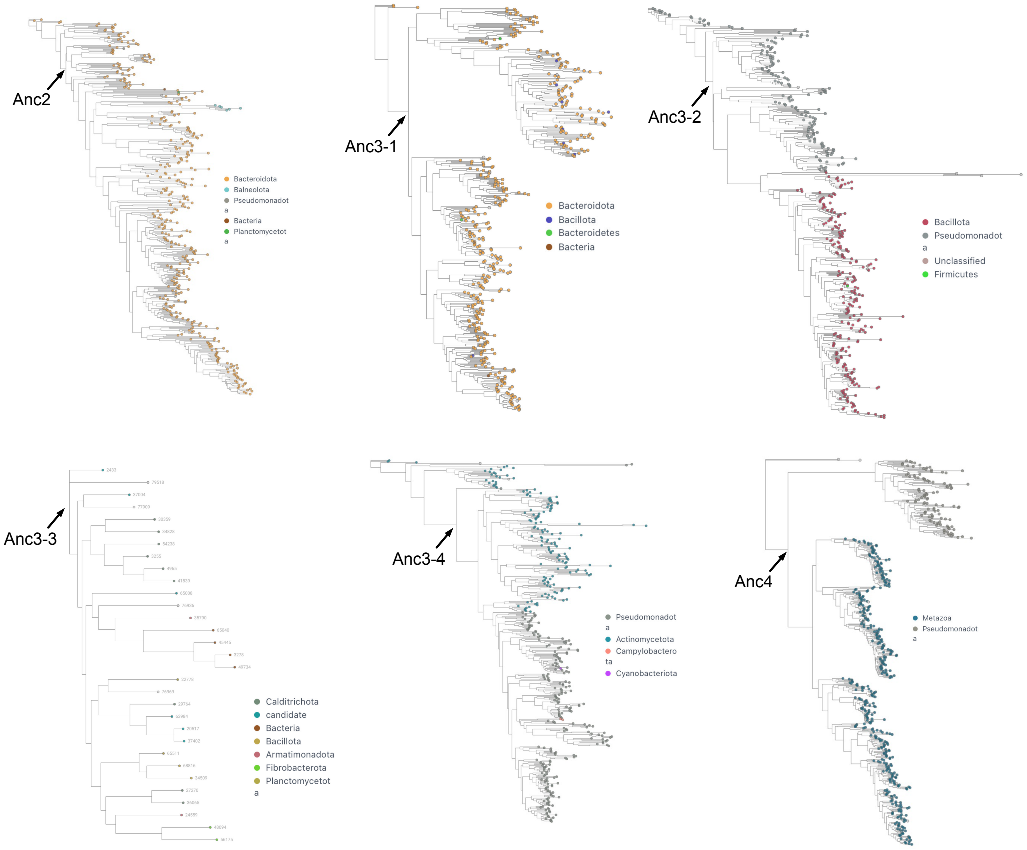


**Figure S11** Ancestral enzyme reconstruction in each of the AP subclusters. Ancestral (Anc) enzymes used in this study are shown by arrows.

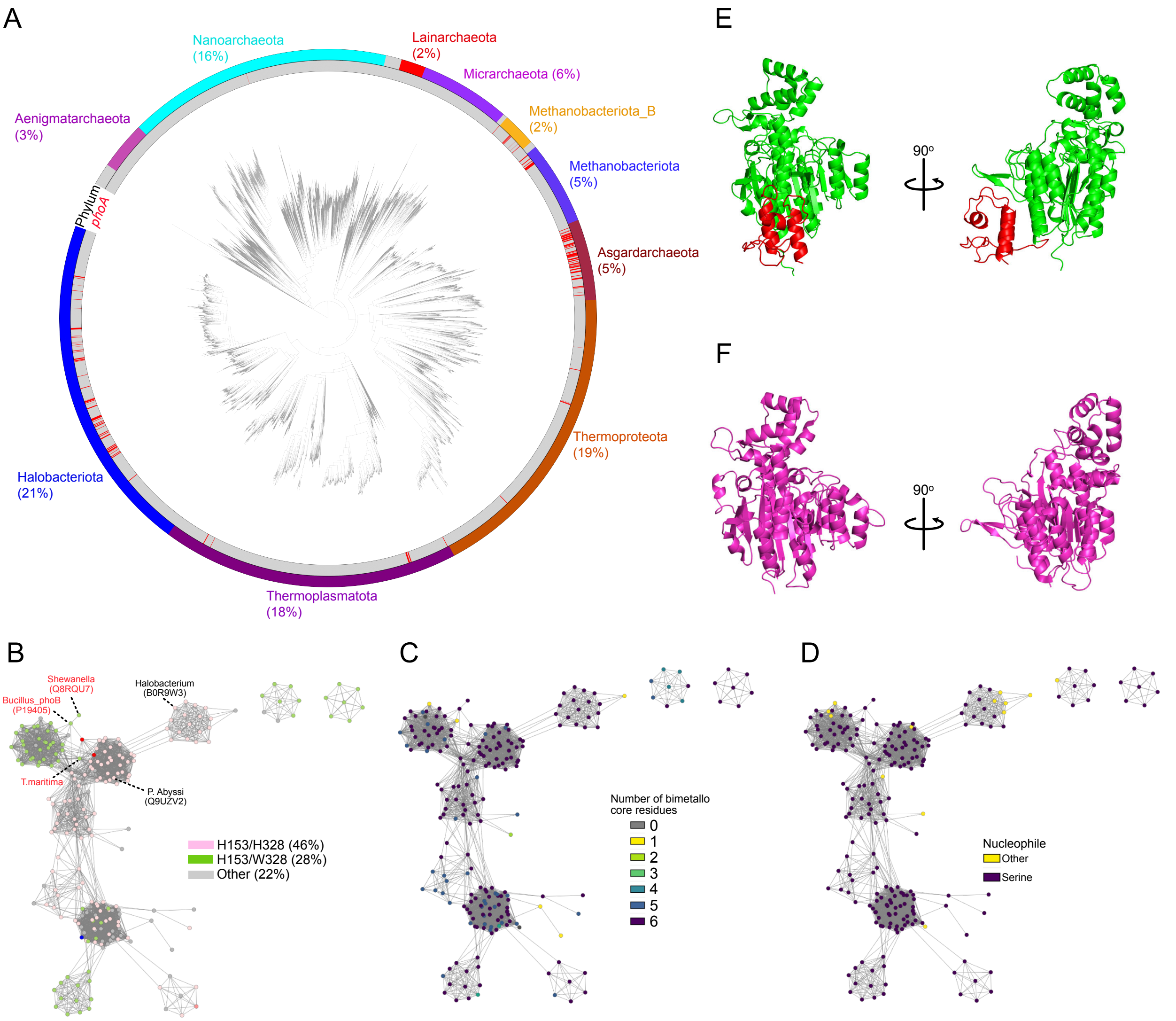


**Figure S12** Phylogenetic tree and SSN analysis of Archaeal APs. (A) The distribution of *phoA* (red) genes mapped onto the Archaeal phylogenetic tree. The lines on the circle indicate the existence of the *phoA* gene. Archaea phyla were also mapped onto the tree. (B-D) Distribution of Archaeal APs was shown with a combination of amino acid residues on 153 and 328 positions (B), number of bimetallo site residues (C), and nucleophile serine (D). Enzymes in red letters indicate bacterial APs with sequences relatively similar to archaeal APs, and those in black letters indicate the archaeal APs used in this experiment. (E), (F) Estimated crystal structure of *P. abyssi* and *H. salinarum* (PDB: 2X98) APs. The C-terminal domain (red domain in *P. abyssi* structure) was removed from the expression vector.

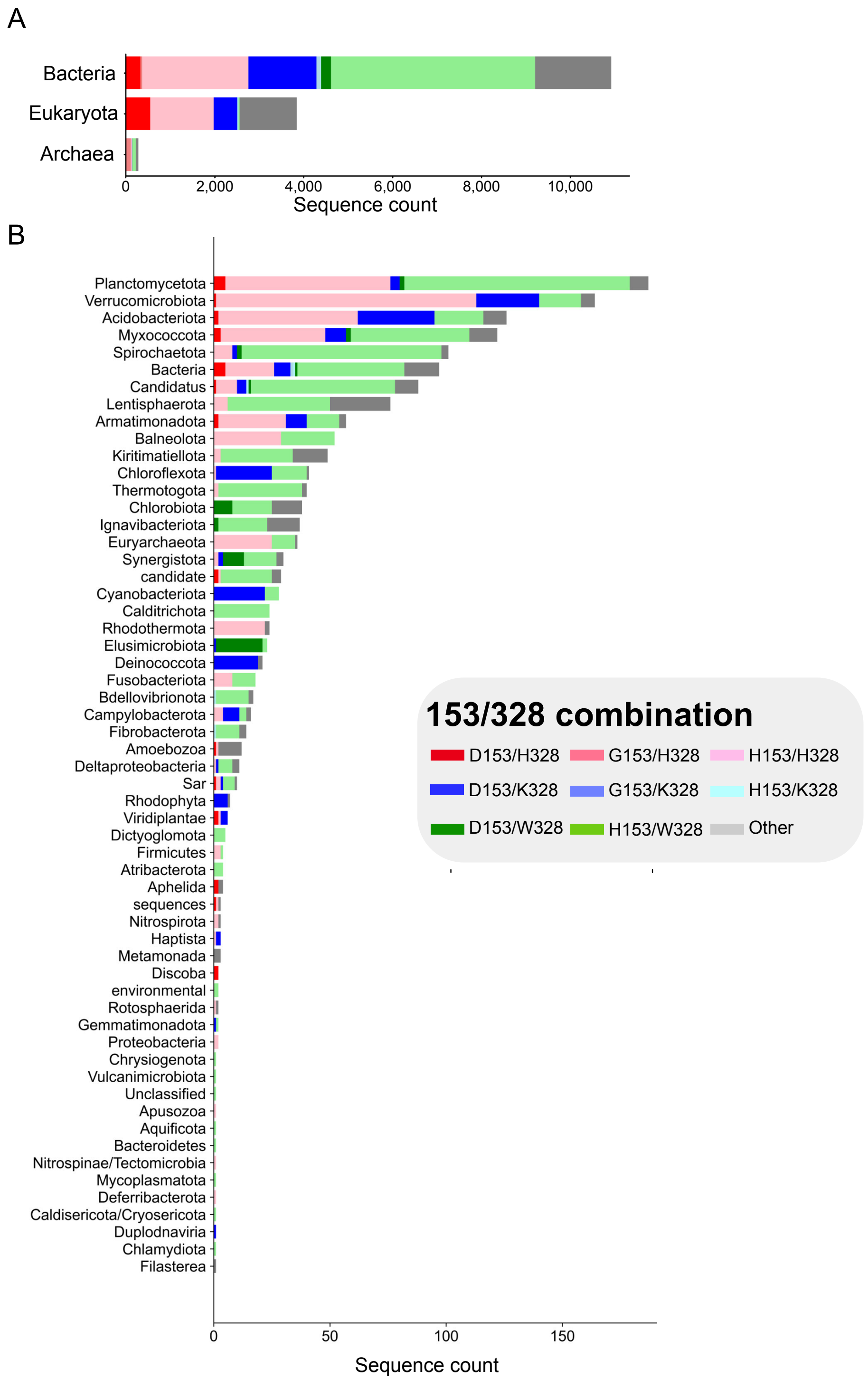


**Figure S13** Selection of the combination of positions 153 and 328 in each domain (A) and phylum (B) within the AP subcluster.

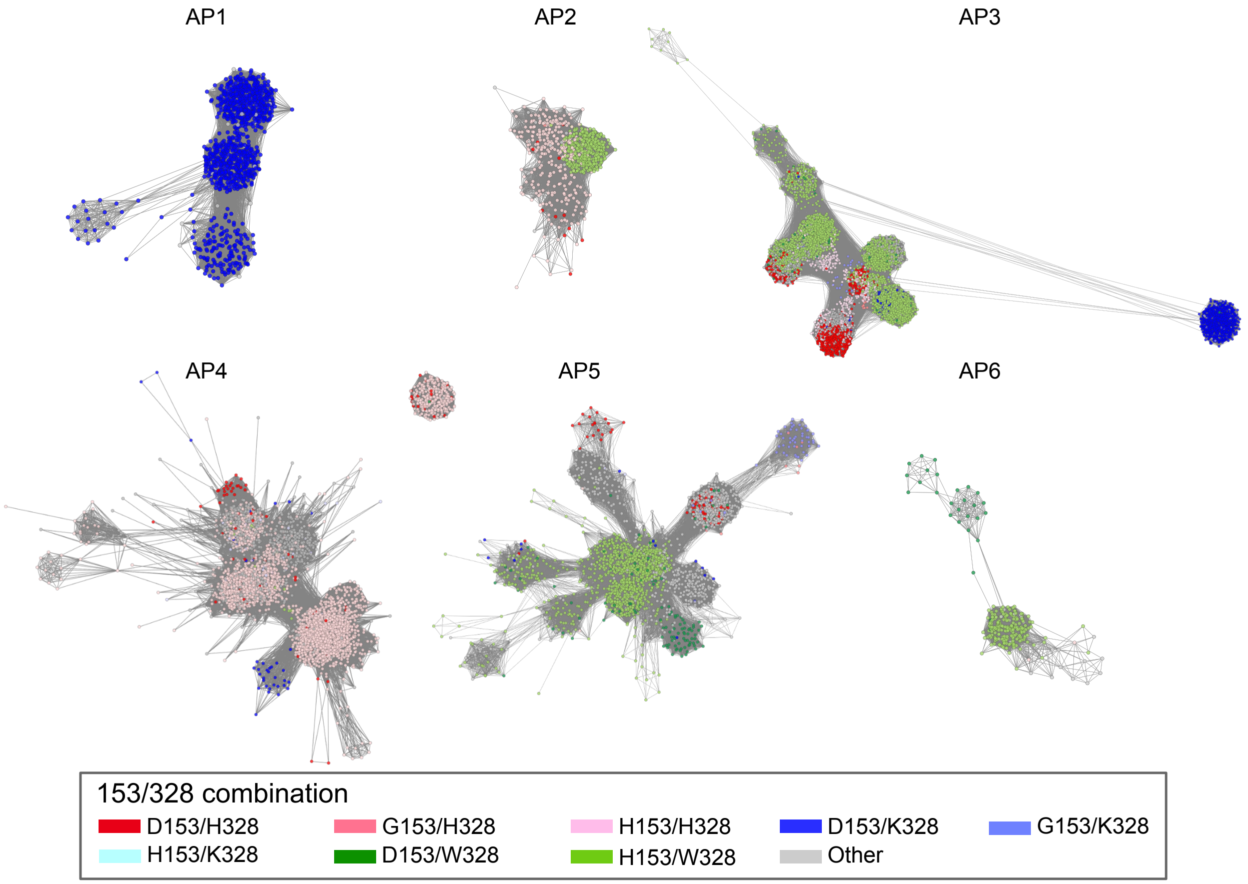


**Figure S14** Mapping positions 153 and 328 combinations on SSN in AP subclusters.

### Materials and Methods

#### Collecting and curating sequence datasets

The sequences of alkaline phosphatase (AP) and its superfamily members were collected from the Pfam database (PF00245, accessed in March 2023), the NCBI Reference Sequence (RefSeq) database (accessed in March 2023), and the Joint Genome Institute (JGI) Integrated Microbial Genomes (IMG) database (accessed in July 2020). To retrieve sequences from the RefSeq and JGI databases, we performed Hidden Markov Model (HMM) searches. A profile HMM (pHMM) for AP was constructed using a multiple sequence alignment (MSA) of several annotated AP sequences generated by ClustalW. AP homologs were identified using an E-value threshold of <1 for the JGI database and <10 for the RefSeq database. For metagenomic APs, redundant sequences were removed using CD-HIT at sequence identity thresholds of 90%, 70%, and 50%. Sequences with lengths outside the target range (<250 or >650 amino acids) were excluded. Pairwise alignments were then performed between the collected metagenomic sequences and *E. coli* AP (*ec*AP). Sequences lacking the bimetallo binding site or the nucleophilic serine residue were removed. The resulting sequences were merged and further clustered using CD-HIT at a 90% identity threshold to reduce redundancy. The final AP superfamily dataset comprised 32,843 sequences.

Dehydrogenase sequences, including the previously characterized PtxD (UniProt ID: O69054), were retrieved from the Pfam database (PF02826, accessed in March 2024), which contained 236,000 sequences. Redundant sequences were removed using CD-HIT with an 80% sequence identity threshold, resulting in a final PtxD dataset comprising 79,615 sequences.

For the PhnJ domain of CP lyase, homologous sequences were initially obtained from the Pfam database (PF06007, containing 5,040 sequences, accessed in April 2024). Additional PhnJ sequences were retrieved from the NCBI RefSeq database via an HMM search. The profile HMM (pHMM) was constructed using a multiple sequence alignment of annotated PhnJ sequences (UniProt IDs: P16688 and Q52987). The HMM search against the RefSeq database was conducted with a bit score cutoff of >500, resulting in a total of 12,588 sequences used to construct the CP lyase dataset.

PstS sequences were collected from the Pfam database (InterPro entry IPR024370, accessed in January 2025) and used to construct the PstS dataset, comprising a total of 81,405 sequences.

**Clustering enzyme family**

An all-versus-all BLAST search was performed across all sequence datasets. To cluster enzyme family members, we employed a meta-sequence similarity network (meta-SSN) analysis using the MetaSSN tool (https://github.com/johnchen93/MetaSSN.git). In the meta-SSN, each node represents a cluster of highly similar sequences, defined by BLAST bit scores exceeding a specified threshold. By gradually increasing the bit score cutoff, members of enzyme families were progressively separated into distinct clusters (**Fig. S1**). The types of enzymes in each subcluster were annotated based on manually reviewed sequences from Swiss-Prot in the UniProt database.

Within each AP subcluster, a standard sequence similarity network (SSN) analysis was conducted, in which each node represents a homologous AP sequence and edges indicate sequence similarity (**Fig. 2A**).

**Identifying the *phoA*, *pafA*, *phnJ*, *ptxD*, and *pstS* genes in the GTDB database**

The *phoA*, *pafA*, *phnJ*, and *pstS* genes were searched in the GTDB database using HMM-based searches. First, multiple sequence alignments (MSAs) were generated for each cluster of the target enzymes—AP1 to AP6 (**Fig. 2A**) and the red nodes in **Fig. S1**—using ClustalW. Profile HMMs (pHMMs) were then constructed from these MSAs. Appropriate bit score cutoffs for the HMM searches in GTDB were empirically determined by performing test searches against known databases. With bit score thresholds of 150, 100, 380, and 50, respectively, 88%, 94%, 98%, and 95% of the sequences retrieved by HMM searches were correctly assigned to AP1–6, PafA, PtxD, and PstS (**Fig. S1A, B,** and **D**) subclusters. PhnJ formed a small and highly conserved protein family, consisting of nearly identical sequences. In the meta-SSN analysis, PhnJ sequences could not be separated even at a bit score cutoff of 500 (**Fig. S1C**); therefore, this threshold was used for the HMM search in GTDB. Finally, we searched for these enzyme genes in the GTDB r207 release (comprising 62,291 representative genomes, one per species) using the established bit score cutoffs.

**Identifying *phoA* genes in the Archaeal database**

To identify *phoA* genes in archaeal genomes, we first retrieved 4,416 representative archaeal genomes (one genome per species) from the GTDB r218 release. HMM-based searches were then performed against this archaeal genome dataset to detect *phoA*, *pafA*, *ptxD*, and *phnJ* genes, using the previously constructed pHMMs and the same bit score cutoff values described above.

**Phylogenetic tree reconstruction of bacterial species**

We utilized a previously reconstructed phylogenetic tree of bacterial species [29]. This tree includes 32,255 representative species across all phyla, each with ≥95% genome completeness.

**Phylogenetic tree reconstruction of AP subclusters**

Multiple sequence alignments (MSAs) were generated for each AP meta-node using ClustalW. The alignments were manually curated in Jalview, where poorly aligned regions and N-terminal signal peptides were trimmed. Signal peptides were predicted using SignalP 5.0. The curated MSAs from all meta-nodes were then concatenated and used to infer maximum likelihood phylogenies with IQ-TREE version 1.6.1. Branch supports were assessed using both ultrafast bootstrap approximation (UFBoot2) and the approximate likelihood ratio test (aLRT), each with 5,000 replicates. IQ-TREE analyses were repeated three times using the same input MSA, yielding highly consistent phylogenetic trees with nearly identical topologies.

**Estimating ancestral states in the phylogenetic tree of the bacterial genomes**

To estimate ancestral states on the bacterial genome phylogenetic tree, we used PastML with the Felsenstein81 model and the marginal posterior probabilities approximation (MPPA) method [29, 34].

**Ancestral sequence reconstruction in the AP subclusters**

We performed ancestral sequence reconstruction using the joint reconstruction method implemented in GRASP [37]. Ancestral enzymes (Anc) were inferred based on the MSAs of AP2, AP3-1, AP3-2, AP3-3, AP3-4, and AP4, which were also used to reconstruct the phylogenetic tree of the AP subcluster (**Fig. 2B**). As the inferred phylogenetic trees often included multiple organisms, we selected the most recent common ancestor shared by different taxa (**Fig. S11**). The active site residues of the inferred ancestral enzymes (**Fig. 4A)**, as determined by joint reconstruction, showed marginal reconstruction probabilities of at least 98%.

**Metagenomic AP and PtxD sequences analysis**

The distributions of AP and PtxD in environmental samples were analyzed using the JGI IMG database. Homologous enzymes were retrieved through HMM searches with the same profile HMMs (pHMMs) and bit score cutoffs applied in the GTDB database search. Sequences shorter than 200 amino acids were excluded from the analysis. The previously obtained RecA data were used as a control [30]. The ecological distribution and relative abundance of AP and PtxD in each environmental sample were analyzed as described in our previous study [30].

**Structural prediction**

Extant and ancestral enzyme structures were predicted using ColabFold v1.5.5.

**Data visualization**

All visualizations and phylogenetic analyses were performed using iTOL and Taxonium. For SSN analysis, Cytoscape was employed. Python packages, including Matplotlib and Seaborn, were used for generating all plots. The structures of the enzymes and their active sites were visualized using PyMOL.

**Cloning/Expression/Purification of homologous APs and PafA**

We used a previously constructed plasmid for expressing and purifying *ec*AP [27]. The *ec*AP sequence was located between its native signal peptide (SP) and StrepII-tag. Other homologous AP sequences were synthesized by Twist Bioscience (South San Francisco, CA, USA) and inserted into the *ec*AP-expressing plasmid. The SP of all other homologous APs was predicted using SignalP-5.0 and removed from the sequence designs. The C-terminal domain of *P. abyssi* AP was also removed from the sequence design (**Fig. S12E**). The 153/328 combination variants of each AP were synthesized via site-directed mutagenesis with these plasmids. For chaperone expression, we used the pGro7 plasmid encoding the GroEL/ES chaperonin. The AP-expressing plasmid and the pGro7 plasmid each contain resistance genes for ampicillin and chloramphenicol, respectively.

AP-expressing plasmids were transformed into Shuffle T7 Express Competent *E. coli* (Δ*phoA*) (New England BioLabs, Ipswich, MA, USA) with or without the chaperone-expressing pGro7 plasmid. The cells were plated on LB agar plates containing 100 μg/mL ampicillin for the AP-expressing plasmid, or 100 μg/mL ampicillin and 34 μg/mL chloramphenicol for both the AP-expressing plasmid and pGro7 plasmid. After overnight culturing from a single colony, 2 mL of the culture solution was transferred to 200 mL of LB medium in a 2 L flask and cultured at 37°C until the OD600 reached 0.4–0.6. Protein expression was induced by adding IPTG (final concentration 100 μM) at 16°C, and the bacteria were cultured for 16–24 hours. For co-expression of AP and GroEL/ES chaperonin, L(+)-Arabinose (final concentration 0.1%) (Thermo Fisher Scientific Inc.) was added after transferring the cells into the flask. Chaperonin co-expression was carried out with insoluble and low-expressing APs, including *pa*AP, *ob*AP, *hb*AP, *bs*AP, *rs*AP, *pb*AP, *sb*AP, *as*AP, and *em*AP. Bacteria were harvested by centrifugation and frozen at -80°C for 1 day. The cells were lysed using a lysis buffer (1:1 mixture of B-PER and 50 mM Tris-HCl (pH 8.0) containing 200 mM NaCl, 10 μM $ZnCl_2$, 1 mM $MgCl_2$, 100 μg/mL lysozyme, 1 μL of benzonase per 30 mL buffer, and 100 μM phenylmethylsulfonyl fluoride (PMSF)). A total of 30 mL of lysis buffer was used per 8 g of bacterial pellet. After incubation for 60 minutes at 22°C, cell debris was pelleted, and the supernatant containing the protein was collected. The cell lysate was passed over a Strep-Tactin column (IBA Lifesciences) pre-equilibrated with a wash buffer (50 mM Tris-HCl (pH 8.0) containing 200 mM NaCl, 10 μM $ZnCl_2$, and 1 mM $MgCl_2$). After washing the column with the wash buffer, the protein was eluted using 50 mM biotin dissolved in the wash buffer. Following elution, the buffer was desalted using a desalting column (Econo-Pac 10DG, BioRad, Hercules, CA, USA) and a storage buffer (10 mM HEPES (pH 8.0) containing 50 mM NaCl, 100 μM $ZnCl_2$, and 1 mM $MgCl_2$). Protein solutions were concentrated using a spin filter (Spin filters 10k, Pall Laboratory, Port Washington, NY, USA).

The expression and purification of PafA were carried out using a method almost identical to that for AP, except that the wash buffer for PafA did not include $MgCl_2$.

**Lysate preparation**

For lysate preparation, we used the same plasmid, bacterial strain, LB medium, and buffers as described in the *Cloning/Expression/Purification of Homologous APs* section. The cells were cultured overnight in 5 mL of LB medium. A 10 μL aliquot of the culture solution was transferred to 1 mL of LB medium in a 96-deep well plate (Avantor, Radnor, PA, USA). The cells were cultured at 37°C for 4 hours, and protein expression was induced by adding IPTG (final concentration 100 μM) at 16°C overnight. For the co-expression of GroEL/ES chaperonin and AP, L(+)-Arabinose (final concentration 0.1%) was added. Bacteria were harvested by centrifugation, and 100 μL of lysis buffer was added to the cell pellet, which was incubated for 1 hour at 22°C. The cell debris was spun down, and the supernatant (lysate) and pellet were collected. The pellet was resuspended in 100 μL of wash buffer. The solubility and expression levels of the enzymes were analyzed by SDS-PAGE. Equal volumes of the pellet and lysate fractions were loaded onto the gel.

**Enzyme linked immunosorbent assay (ELISA) for estimating enzyme concentration**

The purified enzyme concentration was estimated by performing an ELISA. The enzyme concentration of APs was measured using a Nanodrop, and 10 ng of enzyme was loaded onto a MaxiSorp 96-well plate (Thermo Fisher Scientific Inc., Waltham, MA, USA). The plate was incubated overnight at 4°C. After removing the enzyme solution, StartingBlock (Thermo Fisher Scientific Inc.) was added and incubated for 1 hour at room temperature (RT). After removing the StartingBlock, the wells were washed with an ELISA wash buffer (50 mM Tris-HCl, pH 7.5, containing 0.05% Tween 20). A 0.1 μg/mL solution of StrepII-tag antibody-HRP (GenScript, Piscataway, NJ, USA) in the wash buffer was applied to the wells, and the plate was incubated for 1 hour at RT with slow shaking on a 2D rocker.

The antibody solution was removed, and the wells were washed with the wash buffer. Substrate solution (1-Step™ TMB ELISA Substrate Solution, Thermo Fisher Scientific Inc.) was then added to the wells and incubated for 15 minutes at RT. The reaction was stopped by adding 2 M sulfuric acid, and the signal was measured using a plate reader with absorbance at 450 nm (Synergy H1, BioTek, Winooski, VT, USA). Purified *ec*AP was used to prepare a calibration curve for determining the protein concentration.

**Bulk ensemble kinetic assay**

A bulk ensemble kinetic assay was performed in a 384-well plate using a plate reader (Synergy H1). To measure phosphomonoesterase (PME) activity, p-nitrophenyl phosphate (*p*NPP) (MilliporeSigma, Burlington, MA, USA) was used as a substrate. Absorbance was measured at 405 nm, and changes in absorbance were recorded at ten-second intervals at 28°C. The reaction solution was prepared by mixing the enzyme (0.01-1.5 µM) and *p*NPP (final concentration 2 mM) in the reaction buffer on ice, then the mixture was added to the wells. The activity was measured after 1 minute to ensure temperature equilibration. Three independent measurements were taken to calculate the mean and standard deviation (*SD*) of enzyme activity. Enzyme activity was determined from the linear fit of the increase in absorbance over time. The reaction buffer consisted of 1 M DEA buffer (pH 9.5) with 200 µM $ZnCl_2$ and 1 mM $MgCl_2$.

Phosphite (Pt) oxidation activity was measured using 2 mM Pt (Thermo Fisher Scientific Inc.) as a substrate. Pt and enzyme (1.5 µM) were mixed in a PCR tube and incubated for 8 hours at 28°C. After incubation, an equal volume of malachite green phosphate assay solution (MilliporeSigma) was added to the reaction mixture to measure phosphate concentration. Absorbance for color formation was measured at 620 nm. Since the addition of malachite green alters the assay conditions, phosphate concentration was measured at the end-point. Three independent measurements were taken to analyze the mean and *SD* of enzyme activity. The reaction buffer for Pt oxidation contained 50 mM HEPES buffer (pH 8.0), 200 µM $ZnCl_2$, and 1 mM $MgCl_2$.

For assays with the thermophilic *tm*AP and *P. abyssi* AP, PME and Pt oxidation activities were measured at 55°C in the plate reader and at 65°C in a PCR machine, respectively. For *tm*AP, 200 µM $CoCl_2$ was used in place of $ZnCl_2$.

Lysate activity was measured using *p*NPP (final concentration 2 mM) and the DEA buffer described earlier. Lysate activity was determined from the linear fit of the increase in absorbance over time and was corrected based on the OD600 value measured before cell collection after enzyme expression.

**Bacterial growth assay with phosphate and Pt**

We used the same bacterial strains employed for the expression and purification of *ec*AP in the growth assay. Bacteria were streaked onto LB agar plates containing 100 μg/mL ampicillin. After overnight incubation at 37°C, bacterial colonies were collected and transferred to MOPS minimal media lacking a phosphorus source. The growth assay was conducted in a 96-well plate with shaking at 37°C. Bacterial growth was monitored by measuring absorbance at 600 nm every 30 minutes for 72 hours. Bacteria were inoculated into MOPS minimal media containing either 500 μM inorganic phosphate (IP) or phosphite, with a final OD600 of 0.001.
